# Artificial Intelligence uncovers carcinogenic human metabolites

**DOI:** 10.1101/2021.11.20.469412

**Authors:** Aayushi Mittal, Sanjay Kumar Mohanty, Vishakha Gautam, Sakshi Arora, Sheetanshu Saproo, Ria Gupta, Roshan S, Prakriti Garg, Anmol Aggarwal, Padmasini R, Nilesh Kumar Dixit, Vijay Pal Singh, Anurag Mehta, Juhi Tayal, Srivatsava Naidu, Debarka Sengupta, Gaurav Ahuja

## Abstract

The genome of a eukaryotic cell is often vulnerable to both intrinsic and extrinsic threats due to its constant exposure to a myriad of heterogeneous compounds. Despite the availability of innate DNA damage response pathways, some genomic lesions trigger cells for malignant transformation. Accurate prediction of carcinogens is an ever-challenging task due to the limited information about *bona fide* (non)carcinogens. We developed Metabokiller, an ensemble classifier that accurately recognizes carcinogens by quantitatively assessing their electrophilicity as well as their potential to induce proliferation, oxidative stress, genomic instability, alterations in the epigenome, and anti-apoptotic response. Concomitant with the carcinogenicity prediction, Metabokiller is fully interpretable since it reveals the contribution of the aforementioned biochemical properties in imparting carcinogenicity. Metabokiller outperforms existing best-practice methods for carcinogenicity prediction. We used Metabokiller to unravel cells’ endogenous metabolic threats by screening a large pool of human metabolites and predicted a subset of these metabolites that could potentially trigger malignancy in normal cells. To cross-validate Metabokiller predictions, we performed a range of functional assays using *Saccharomyces cerevisiae* and human cells with two Metabokiller-flagged human metabolites namely 4-Nitrocatechol and 3,4-Dihydroxyphenylacetic acid and observed high synergy between Metabokiller predictions and experimental validations.

## INTRODUCTION

Eukaryotic cells encounter a large number of structurally and functionally distinct compounds in their life cycle^1–3^. Such interactions often trigger dysregulation of cellular processes, resulting in loss of cellular homeostasis^1,2,4^. Some of these compounds alter the critical components of a cell, such as its genome or the surveillance mechanisms that ensure its integrity, leading the cells to malignant transformation^4–7^. These compounds are called carcinogens^8^. Due to the ever-increasing incidence of cancer worldwide, carcinogens are emerging as a major health hazard^9^. Notably, only 5-10% of the cancer types are heritable in nature, whereas, the majority of the cancers are caused due to exposure to carcinogens^10^. Mechanistic insights about their mode of action is still under exploration, however, multiple studies have revealed that carcinogens also impact the genetic material (DNA) of a cell and globally alter the key cellular machinery such as its epigenome, transcriptome, proteome, and metabolome^4,11,12^. Recent reports suggest that a chemical carcinogen could induce carcinogenicity by directly impairing the epigenome, interfering with the DNA damage response pathway, and/or activating the anti-apoptotic pathways^4,11,12^. It has also been observed that carcinogens trigger alterations in the cellular microenvironment that lead to energy metabolic dysfunction, thereby inducing carcinogenesis^13–15^. These findings suggest complexities involved in carcinogens’ mode of action. Present approaches to zero in on the carcinogens are contingent on expensive (up to $4 million per compound) and time-consuming (more than two years) animal model-based approaches^16,17^. Importantly, these rigorous testing protocols are still questionable for evaluating carcinogenic threats to humans^18^. Strikingly, an average validation involves harvesting of ∼800 animals, raising ethical concerns^16,17^. While *in vivo* experiments offer indisputable identification of carcinogens, Artificial Intelligence can significantly accelerate pre-screening of the ever-expanding space of compounds that includes new drugs, chemicals, and industrial by-products^19–22^. To date, numerous computational methods have been proposed for carcinogenicity prediction. In most cases, quantitative structure-activity relationship (QSAR) models have been used for the classification of carcinogens and non-carcinogens^21–28^. Structural alert-based expert systems have also been proposed where the carcinogenic potential of a compound is estimated using two-dimensional structural similarities^29,30^. QSAR models offer decent predictive performance due to their ability to detect specific functional groups that are reported or linked to carcinogenicity. Some of the prominent functional groups include nitro compounds, aromatic amines, polycyclic aromatic hydrocarbons, and polychlorinated biphenyls^24,25,31,32^. To date, all predictive models for carcinogenicity prediction are supervisory in nature and use a limited number of known carcinogens and non-carcinogens for model training. Moreover, a large number of these prediction models heavily rely on genotoxicity (or mutagenicity) alone, leading to a suboptimal predictive performance on non- or mildly genotoxic carcinogens. Although there is a strong link between genotoxicity and carcinogenicity, genotoxicity alone is insufficient to explain the mode of action of all carcinogens. Some classical examples of non-genotoxic carcinogens include 1,4-dichlorobenzene (tumor promoter)^33^, 17β-estradiol (endocrine-modifier)^34^, 2,3,7,8-tetrachlorodibenzo-p-dioxin (receptor-mediator)^35^, and cyclosporine (immunosuppressant)^36^. On the other hand, N-Nitroso-2-hydroxy morpholine is genotoxic but non-carcinogen^37,38^. Given the complexity of the chemical space, linear approaches relying either on a limited training dataset or measurement of genotoxic/mutagenic properties fail to generalize on unseen compounds with discordant structural properties.

With the advancement in functional assays, it has been established that a potential carcinogen might induce cellular proliferation^39,40^, genomic instability, oxidative stress response, anti-apoptotic response^40^, and epigenetic alterations^41^. Additionally, a large number of carcinogens are known to be electrophilic in nature^42–44^. We developed Metabokiller, a method that utilizes an ensemble classification approach that harnesses the aforementioned biochemical properties. Metabokiller identified a number of human metabolites that might possess carcinogenic properties. We selected two previously uncharacterized human metabolites i.e. 4-Nitrocatechol and 3,4-Dihydroxyphenylacetic acid, which were predicted by Metabokiller as potential carcinogens and experimentally validated their predicted carcinogenicity using an array of functional assays and deep RNA sequencing.

## RESULTS

### Metabokiller: Biochemical properties-driven Ensemble Model for Carcinogenicity prediction

Cancer cells are highly proliferative, and possess altered epigenetic signatures, elevated reactive oxygen species (ROS) levels, and activated anti-apoptotic pathways^45^. Moreover, it has been observed that carcinogens possess electrophilicity, and therefore, might pose a threat to genomic stability^42,43^. Unlike other QSAR-based models for carcinogenicity prediction that largely use limited available information about the experimentally-validated carcinogens and non-carcinogens, Metabokiller tracks a set of well-known, carcinogen-centric biochemical properties. Models created for the individual properties are finally combined into an ensemble model that evaluates the query compound for these biochemical properties and gives a consensus score indicating the carcinogenicity potential **(Figure 1a, Supplementary Figure 1a)**. To build Metabokiller, we first manually curated and compiled datasets containing information about compounds that are reported to impact cellular proliferation, genomic stability, oxidative state, epigenetic landscape, and apoptotic response. We also compiled a dataset comprising information about *bona fide* electrophiles and non-electrophiles. Each of the six independent datasets (electrophilic properties, epigenetic modifications, genomic instability, oxidative stress, proliferative properties, and anti-apoptotic properties) is segregated into pro-(Class 1) or anti-/no- (Class 0) activity categories, collectively containing 35,668 compounds (combined data annotated as MK_Tn_) **(Figure 1b, Supplementary Table 1)**. Next, to gain deeper insights into the chemical heterogeneity between the classes, we performed Principal Component Analysis (PCA) using the Extended-connectivity fingerprints (ECFPs) as features and observed a high degree of chemical heterogeneity in almost all the cases. Inter-class functional group comparison revealed the selective enrichment or de-enrichment of distinct functional groups in every dataset. Collectively, these results indicate a higher degree of functional and chemical heterogeneity between the classes for each of these biochemical properties (**Figure 1c**). We next built six independent classification models featuring the aforementioned biochemical properties. To obtain the best performing models, we tried three different feature extraction methods i.e. bioactivity-based descriptors (Signaturizer library)^46^, chemistry-based molecular descriptors (Mordred software)^47^, and graph-based features (DeepChem library)^48^. In addition to these diversified features, we also tried multiple machine learning/deep learning-based classification algorithms for model building such as Random Forest (RF), Multilayer Perceptron (MLP), k-Nearest Neighbor (KNN), Support Vector Machine (SVM), Stochastic Gradient Descent (SGD), Logistic Regression (LR), GraphConvModel (GCM), Attentive FP (AFP), Graph Convolution Network (GCN), and Graph Attention Network (GAT) **(Supplementary Figure 1b)**. The generic workflow for the model building includes random splitting of the compiled biochemical datasets into training and testing data, feature extraction from the compound SMILES using three orthogonal methods (Mordred, Signaturizer, and graph-based), selection of the highly variable, least correlated, and more important features using Boruta^49^, handling class imbalance using Synthetic Minority Oversampling Technique (SMOTE)^50^, implementation of an array of classifiers for model building, and testing of all models on the same unseen testing dataset. We uniformly performed these steps to build twelve different classification models for each biochemical property and subsequently selected the best performing and most stable models i.e. KNN (anti-apoptotic), MLP (electrophiles), SVM (epigenetic modifications), RF (genomic instability), MLP (oxidative stress), and RF (proliferation) **(Supplementary Figure 1b, Supplementary Figure 2a)**. Notably, among all the tested conditions, Signaturizer-based models outcompeted other models. We next optimized these base models further by performing randomized-search-based hyperparameter tuning **(Supplementary Table 4)** and performed rigorous testing of the selected parameters using bootstrapping (twenty repetitions) and 10-fold cross-validation techniques **(Figure 1d-g, Supplementary Figure 1a, Supplementary Figure 2b)**.

**Figure 1:**
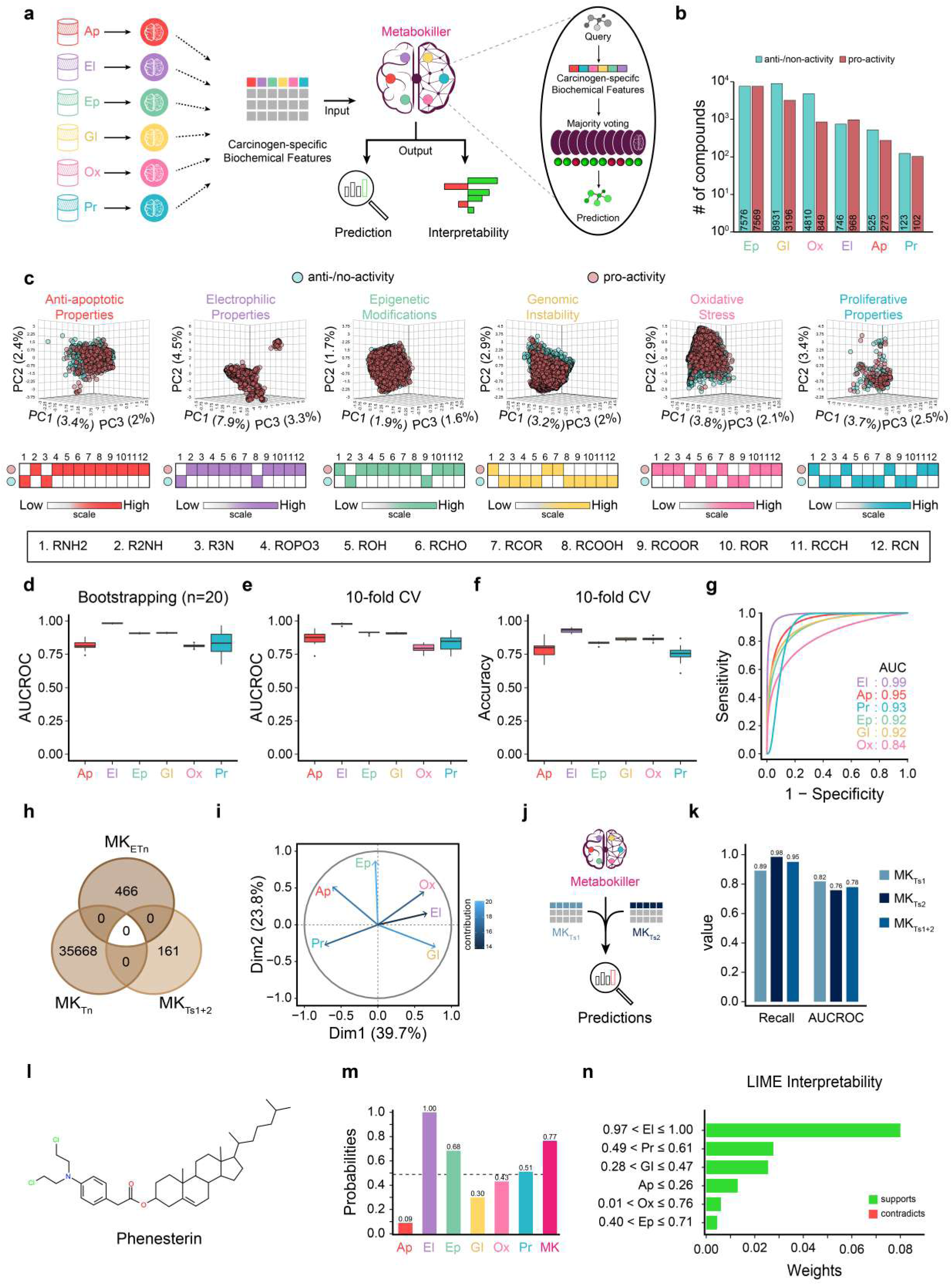
Metabokiller, an explainable artificial intelligence-driven ensemble model for carcinogenicity prediction. **(a)** Schematic representation depicting the key steps involved in building Metabokiller. Of note, six independent classification models that form the core of Metabokiller prediction engines are highlighted in different colors. Prediction probabilities from these six models are used as carcinogen-specific biochemical features for ensemble models, and the final prediction of the query molecule is performed based on the majority voting method. Metabokiller offers two main modules, i.e. prediction and interpretability. **(b)** Bar graph depicting the number of compounds used in the training datasets of the six indicated biochemical models. Of note, turquoise and indian-red bars indicate the number of compounds assigned as anti-/no- and pro-activities in the indicated conditions, respectively. **(c)** Principal Component Analysis revealing the chemical heterogeneity between the two classes (pro- and anti-/no-) of the indicated datasets. Heatmaps at the bottom depicting the relative enrichment of the indicated functional groups in both classes. **(d)** Box plot depicting the AUCROC values of the bootstrapping (20 repetitions) of the indicated models. **(e)** Box plot depicting the AUCROC values of the 10-fold cross-validation of the indicated models. **(f)** Box plot depicting the accuracy values of the 10-fold cross-validation of the indicated models. **(g)** AUC (Area under the curve) plots representing the performance of the best models obtained using 10-fold cross-validation of the indicated best models. **(h)** Venn diagram depicting no overlaps between the compounds constituting six independent biochemical property-based models (annotated as MK_Tn_), training dataset used to build the Metabokiller ensemble model (annotated as MK_ETn_), and the external testing data containing information about experimentally-validated carcinogens and non-carcinogens (annotated as MK_Ts1+2_). **(i)** Variables factor map (PCA) depicting the direction and contribution of all the six variables (individual models) representing the experimentally-validated carcinogens (MK_ETn_) in the Eigenspace. **(j)** Schematic representation of the projection of the external testing datasets containing information about experimentally-validated carcinogens and non-carcinogens (annotated as MK_Ts1_ and MK_Ts2_) on Metabokiller. **(k)** Bar graph depicting the AUCROC values of the Metabokiller performance on external testing datasets annotated as MK_Ts1_, MK_Ts2,_ and MK_Ts1+2_). **(l)** Chemical structure of Phenesterin, a known chemical carcinogen. **(m)** Bar graph depicting the prediction probabilities of the indicated models for Phenesterin. **(n)** Bar graph representing the results of Local interpretable model-agnostic explanations (LIME), an interpretability module of Metabokiller, highlighting the key biochemical properties of Phenesterin responsible for carcinogenicity.

We finally built an ensemble model (Metabokiller) by leveraging the prediction probability outputs from these six models under a unified machine learning-based framework for carcinogenicity prediction. Methodologically, we took experimentally validated *bona fide* carcinogens and non-carcinogens (annotated as MK_ETn_) **(Supplementary Table 2)** and computed their carcinogen-specific biochemical properties using our six biochemical properties-based models. Next, using these prediction probabilities as features, we built twenty distinct Gradient Boosting Machines (GBM)-based models by using bootstrapping technique **(Supplementary Figure 2c)**. This ensemble model uses the majority voting method for carcinogenicity prediction of the query compounds **(Figure 1a, Supplementary Figure 1a)**. Of note, the MK_ETn_ dataset (# of compounds = 466) was collected from ISSCAN^51^ and CCRIS^52^ databases and had no overlaps with MK_Tn_, therefore, avoiding any dataset-dependent biases **(Figure 1h, Supplementary Figure 2d)**. Moreover, carcinogens in this dataset harbor non-redundant, and diversified biochemical properties as indicated by the vectors in the Eigenspace (**Figure 1i**). Notably, we tested six classifiers for building an ensemble model (Metabokiller) and selected the GBM classifier that harbored high precision and recall values, essential parameters for carcinogenicity prediction **(Supplementary Table 3)**.

Finally, to evaluate the performance of Metabokiller on unseen external datasets, we first compiled a list of experimentally-validated carcinogens and non-carcinogens from varied sources^51–54^ (annotated as MK_Ts1_ and MK_Ts2_) **(Figure 1j, Supplementary Table 2)**. Of note, these external datasets contained a high degree of chemical heterogeneity as shown in the Principal Component Analysis (PCA) and in the differential enrichment analyses of the functional groups **(Supplementary Figure 2e)**. We next tested these datasets on Metabokiller and attained highly competitive performance scores. Notably, the Recall of Metabokiller on the unseen datasets is substantially high, suggesting its robustness in classifying carcinogens (true positives) which is extremely vital for the prediction models in the healthcare domain **(Figure 1k, Supplementary Figure 2f)**. In summary, our results suggest that in contrast to all the previous methods, Metabokiller provides an orthogonal means for carcinogenicity prediction, and exhibits higher prediction performance on unseen data. In addition to its strikingly high performance, the biochemical property-focused Metabokiller, by the virtue of its construction, offers interpretability. Moreover, we also implemented Local interpretable model-agnostic explanations (LIME) algorithm^55^ that imparts interpretability features to Metabokiller (**Figure 1l-n**). Further details of the approach can be found in the Methods section.

### Metabokiller outperformed existing methods for Carcinogenicity prediction and unfolded endogenous metabolic threats

Next, we performed a comparative analysis using Metabokiller alongside nine other widely accepted tools or methods for carcinogenicity predictions, namely, Protox-II^22^ (annotated as P-II), CarcinoPred-EL (annotated as C-RF (Random Forest), C-SVM (Support Vector Machine), and C-XGB (XG-Boost))^21^, lazar (annotated as L-rt (rat), L-ro (rodents), and L-mo (mouse))^56,57^, and versatile variable nearest neighbor (vNN-r (restricted), and vNN-nr (non-restricted))^58^ (**Figure 2b**). Of note, unlike Metabokiller, all these methods solely rely on the limited available information of (non)carcinogens (Protox-II, lazar, and CarcinoPred-EL) and (non)mutagens (vNN) **(Supplementary Figure 3a)**. For comparative analysis, we first collated a list of experimentally-validated carcinogens and non-carcinogens and used them as an independent dataset (I.D.) **(Supplementary Figure 2d, Supplementary Table 2)**. Of note, I.D. does not contain any overlapping compound with datasets used for training or testing the Metabokiller models (MK_Tn_, MK_ETn_, and MK_Ts1+2_), or in the training dataset of other models used for cross-comparison (**Figure 2a**). Comparative analyses revealed the superior performance of Metabokiller as compared to all other methods in most performance metrics (**Figure 2d-h**). As discussed earlier, Metabokiller attains the highest Recall (value=0.98; max value=1) allowing the accurate predictions of true positives (carcinogens), whereas all other methods severely failed in this parameter (ranging from 0.00 to 0.72; max value=1) (**Figure 2f**). Moreover, Metabokiller also attained the highest overall accuracy level (value=0.81; max value=1), in contrast to other methods (ranging from 0.30 to 0.70; max value=1) (**Figure 2e**). Notably, except for Metabokiller, Protox-II, and vNN-nr, all other tested methods made predictions only on a subset of query molecules, further reflecting their deficiencies in predicting all query compounds (**Figure 2c**). To gain deeper insights into the predicted results, we next calculated and compared the number of true positive, true negative, false-positive and false-negative predictions across all methods. We observed a substantial decline in TP/FN ratio in all other methods in contrast to Metabokiller **(Supplementary Figure 3b)**. These results collectively indicate that the selected biochemical properties leveraged by Metabokiller holistically capture the chemical and functional properties of carcinogens, thereby, enabling their accurate classification.

**Figure 2:**
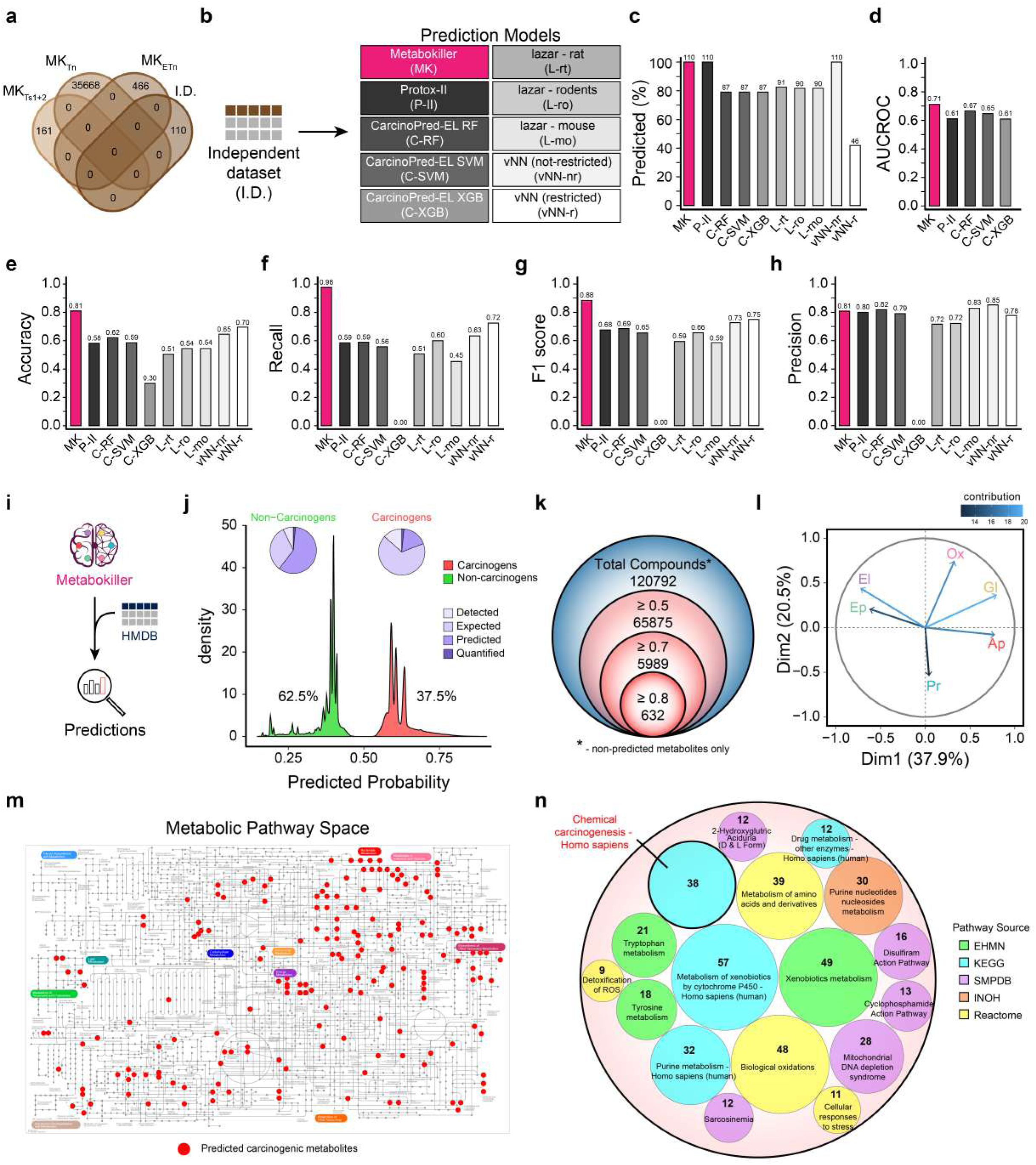
Metabokiller predicted human metabolites possessing carcinogenic potential, and outperformed other prediction methods. **(a)** Dataset used to evaluate and cross-compare the performance of Metabokiller with widely accepted models/tools contains no compound overlaps, represented using Venn Diagram. The datasets used to build the six independent biochemical properties-based models are annotated as MK_Tn_, the training dataset used to build the Metabokiller ensemble model is annotated as MK_ETn_, the external testing data containing information about experimentally-validated carcinogens and non-carcinogens is annotated as MK_Ts1+2_, and the dataset selected for the cross model performance evaluation is annotated as Independent Dataset (I.D.). **(b)** Diagrammatic representation of the experimental workflow to cross-compare the performance of Metabokiller alongside indicated methods for carcinogenicity predictions. These methods include Protox-II (annotated as P-II), CarcinoPred-EL (annotated as C-RF (Random Forest), C-SVM (Support Vector Machine), and C-XGB (XG-Boost)), lazar (annotated as L-rt (rat), L-ro (rodents), and L-mo (mouse)), and versatile variable nearest neighbor (vNN-r (restricted) and vNN-nr (non-restricted)). **(c)** Percentage bar graph depicting the number of compounds predicted by the indicated methods/models. **(d)** Bar graph depicting the AUCROC values of the indicated models on Independent Dataset (I.D.). **(e)** Bar graph depicting the Accuracy values of the indicated models on Independent Dataset (I.D.). **(f)** Bar graph depicting the Recall values of the indicated models on Independent Dataset (I.D.). **(g)** Bar graph depicting the F1 score of the indicated models on Independent Dataset (I.D.). **(h)** Bar graph depicting the Precision values of the indicated models on Independent Dataset (I.D.). **(i)** Schematic representation depicting the testing of Human Metabolome Database (HMDB) metabolites on Metabokiller. **(j)** Overlapping density plots depicting the distribution of prediction probabilities of human metabolites obtained using Metabokiller. Of note, metabolites predicted as carcinogens and non-carcinogens are represented in red and green color, respectively. Pie charts at the top represent the distribution of metabolites based on their annotated status in HMDB i.e., detected, expected, predicted, and quantified. **(k)** Venn diagram depicting the number of predicted-carcinogenic human metabolites, further segregated based on their prediction probability cutoffs. **(l)** Variables factor map (PCA) depicting the direction and contribution of all the six variables (individual models) representing the predicted carcinogenic metabolites from HMDB in the Eigenspace (probability cutoff ≥ 0.5). **(m)** Projection of the predicted carcinogens (indicated as red dots; probability cutoff ≥ 0.7) on the human metabolic space, achieved using iPath Web Server^94^. **(j)** Circle packing chart representing metabolic pathway enrichment results of the predicted carcinogenic metabolites (probability cutoff ≥ 0.7) using Metabokiller. Of note, the pathway information is gathered from The Edinburgh Human Metabolic Network (EHMN), The Integrating Network Objects with Hierarchies (INOH), Reactome, Kyoto Encyclopedia of Genes and Genomes (KEGG), and Small Molecule Pathway Database (SMPDB), and the analysis was performed using Integrated Molecular Pathway Level Analysis (IMPaLA)^95^.

We next aimed to predict the cellular metabolites that harbor carcinogenic properties by leveraging Metabokiller. To achieve this, we performed a large-scale *in silico* screening of the human metabolome. Methodologically, we projected all human metabolites cataloged in the Human Metabolome Database (HMDB)^59^ on Metabokiller and calculated their carcinogenicity probabilities (**Figure 2i**). HMDB contains information about the small molecule metabolites found in the human body, accounting for a total of 2,17,921 metabolites. These metabolites were further classified into four groups based on their status i.e. detected, expected, predicted, and quantified. Interestingly, about 37.5% of the tested metabolites were predicted to possess carcinogenic properties (probability cutoff ≥ 0.5) (**Figure 2j**). Since HMDB also contains information about predicted metabolites, for the downstream analysis we only focused on non-predicted metabolites. At stringent cutoffs of ≥ 0.7 **(Supplementary Table 5)** and ≥ 0.8, we obtained 5989 and 632 predicted carcinogenic metabolites, respectively (**Figure 2k**). In order to determine the interrelationship between the individual biochemical properties represented in predicted carcinogenic metabolites, we computed and compared their prediction probability vectors in the Eigenspace. Our results indicate that almost all the six biochemical properties comprehensively, and non-redundantly capture the carcinogenic feature space, indicating the robustness of our biochemical-based assay in predicting carcinogenicity (**Figure 2l**). Next, to dissect the molecular pathways in which these predicted carcinogenic metabolites are involved, we projected them on the metabolic pathway space and observed their enrichment in distinct metabolic pathways (**Figure 2m**). Moreover, to gain deeper functional insights, we also performed over-representation analysis on the predicted carcinogenic metabolites. Functional analysis revealed that the predicted carcinogenic metabolites (probability cutoff ≥ 0.7) are specifically associated with tyrosine and tryptophan metabolism, nucleotide metabolism, metabolic pathways associated with drug metabolism, xenobiotics, oxidative stress, etc (**Figure 2n**). Notably, our unbiased functional analysis also captured 38 previously known carcinogenic metabolites. Some of the potential endogenous carcinogenic metabolites include Melphalan^60^, Sulfur mustard^61^, 1,2-Dichloroethane^62^, chloral, Thiotepa, N-nitrosodimethylamine (NDMA)^63^, 2,2,2-Trichloroethanol^64^ and others. As an alternative case study, we also used Metabokiller and its inherent interpretability module to decode the functional relevance of oncometabolites in tumor biology. We first reanalyzed the recently integrated cancer metabolomics dataset^65^, and performed differential enrichment analysis to identify tumor enriched or de-enriched metabolites by computing log_2_ fold change. We then projected all tumor-associated metabolites on Metabokiller **(Supplementary Figure 3c)** and computed a dataset-specific correlation between the log_2_ fold change (p-value < 0.05) and the carcinogenicity prediction probabilities. In most cases, we observed a weak negative correlation, suggesting de-enrichment of carcinogenic metabolites **(Supplementary Figure 3d)**. Of note, an in-depth investigation of the enriched metabolites (log_2_ fold change ≥ 1 and p-value < 0.05) identified some of the already reported carcinogenic metabolites, which have been earlier implicated as oncometabolites. These include 2-hydroxyglutarate, succinate, fumarate, N-acetylaspartate, adenylosuccinate, polyamines (putrescine, spermidine, and spermine), etc^66–70^ **(Supplementary Figure 3e, f)**. In summary, by using an orthogonal approach leveraging both the biological and chemical properties of carcinogens, Metabokiller revealed a subset of human metabolites that possess carcinogenic properties.

### Experimental validation of Metabokiller predictions on two previously uncharacterized potential carcinogenic metabolites

Large-scale *in silico* screening of human metabolomes led to the identification of multiple previously uncharacterized carcinogenic metabolites. Next, to validate the Metabokiller predictions, we selected two previously uncharacterized human metabolites i.e., 4-Nitrocatechol (4NC) and 3,4-Dihydroxyphenylacetic acid (DP). Of note, DP is a metabolic intermediate of the tyrosine metabolism pathway, whereas 4NC is involved in aminobenzoate degradation **(Supplementary Figure 4a)**. Importantly DP was predicted to be a non-carcinogen by all other methods except Metabokiller, while there were conflicting results for 4NC **(Supplementary Table 6)**. Metabokiller predicted DP as a potential carcinogen with a prediction probability of 0.71. Moreover, evaluation of the individual models’ probabilities revealed that carcinogenic properties of DP are due to its high genotoxic potential, its electrophilic nature, its ability to alter the epigenetic landscape, and having pro-proliferative and anti-apoptotic properties. Of note, in the case of oxidative stress, we observed below threshold values (cutoff ≥ 0.5) for the DP, suggesting its inability to induce reactive oxygen species (oxidative stress). Similar to DP, 4NC was also predicted as a potential carcinogen with almost equivalent prediction probability (0.70), however, unlike DP, 4NC qualifies in all the individual models, except for epigenetic alteration property (**Figure 3a**). We first investigated the potential of both of these metabolites to induce genotoxicity/genomic instability by performing a single-cell gel electrophoresis-based comet assay, a standard method used to measure DNA breaks and lesions^71^. These experiments were performed on *Saccharomyces cerevisiae* (budding yeast). Briefly, the yeast cells were exposed to these metabolites at different concentrations, and post-exposure, the lysed cells were stained using Giemsa dye. We used hydroxyurea (HU), a known DNA repair inhibitor with mutagenic and genotoxic abilities as a positive control in these experiments^72^. Quantitative assessment of the comet tail lengths revealed a dose-dependent statistically significant increase in the case of 4NC and DP treatment, suggesting their potential to induce genomic instability (**Figure 3b**). Since the training dataset for genomic instability also contains compounds screened for the AMES test, we next asked whether 4NC and DP also possess mutagenic/genotoxic properties. To address this hypothesis, we performed mutagen testing experiments using the high-throughput mutagenesis assay and screened for canavanine-resistant mutants. Of note, we tested different concentrations of 4NC and DP alongside negative vehicle controls. Our results revealed a significant increase in the frequency of canavanine mutants in the treated conditions in contrast to the negative control as indicated by the bean plot. These results validate the mutagenic/genotoxic properties of both DP and 4NC (**Figure 3c**). Higher concentrations of certain biomolecules often cause pleiotropic effects and indirectly impair the key cellular processes. To test whether the concentrations of the metabolites used in mutagenesis assays significantly alter the survival and physiology of the budding yeast, we performed a propidium iodide-based cell viability assay, and observed no profound cell deaths in all the tested concentrations **(Supplementary Figure 4b, c)**. We next wondered whether the observed genotoxicity effect of 4NC is due to the induction of oxidative stress. Notably, 4NC, but not DP were predicted by Metabokiller to induce reactive oxygen species (ROS) levels. We next tested these metabolites for their potential to induce oxidative stress via a fluorometric assay using 2’,7’-dichlorofluorescein diacetate (DCFH-DA) dye. Our results suggest a selective increase in the ROS levels in the case of 4NC, but not DP, at all tested concentrations, suggesting a larger synergy with the prediction probabilities of Metabokiller **(Figure 3d, Supplementary Figure 4d)**. In order to establish a direct link between the 4NC-mediated elevated ROS levels and genotoxicity, we performed high-throughput mutagenesis rescue experiments by using a ROS scavenger. We observed a significant reduction in the number of canavanine resistant mutants in cells co-incubated with the ROS scavenger suggesting ROS-induced genotoxicity (**Figure 3e**).

**Figure 3:**
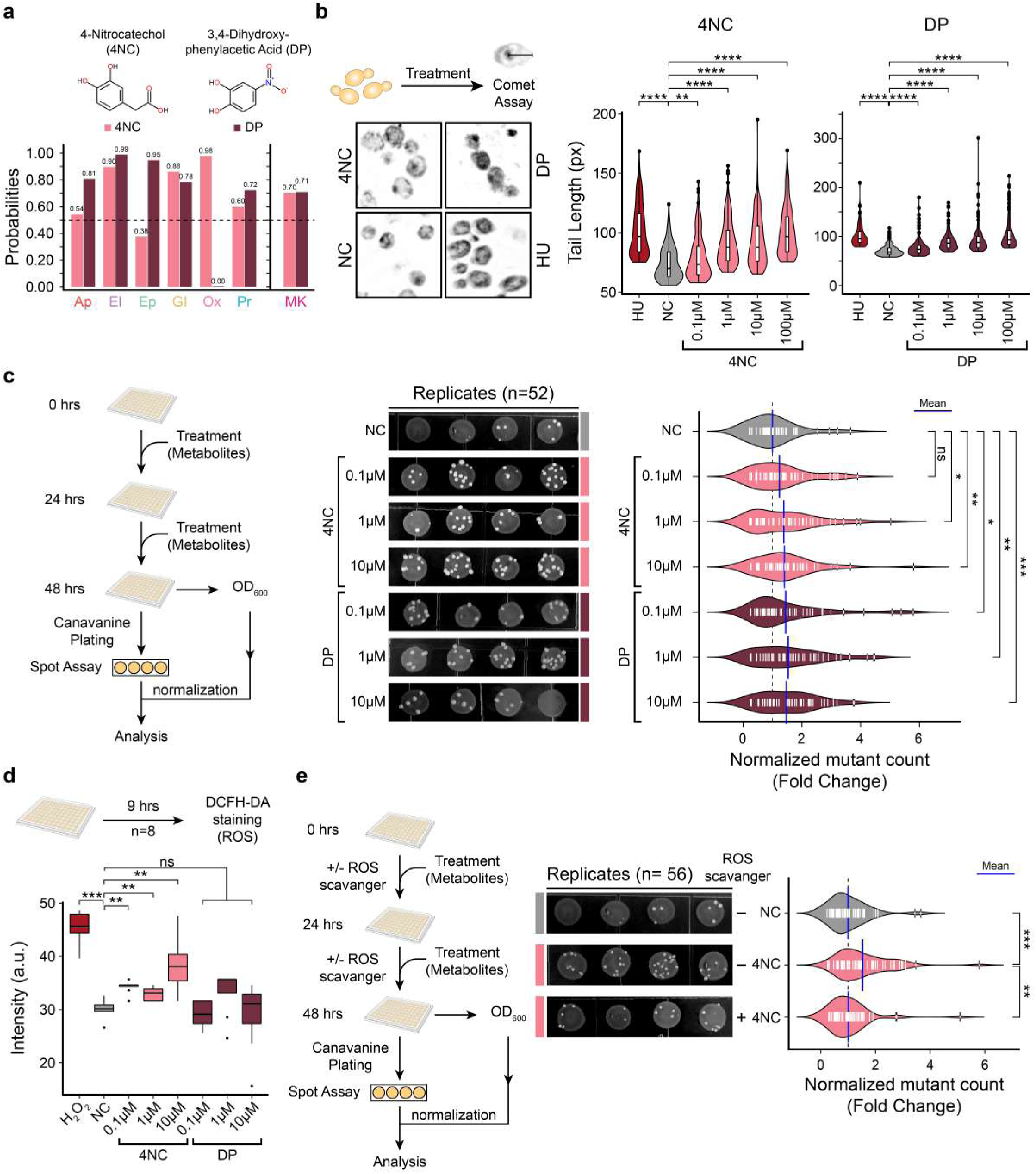
High-throughput assays experimentally validate the Metabokiller predictions. **(a)** Bar graph denoting the prediction probabilities of the six biochemical properties and the ensemble model (Metabokiller) for the two previously uncharacterized human metabolites i.e. 4-Nitrocatechol (4NC), and 3,4-Dihydroxyphenylacetic acid (DP). Of note, the chemical structures of the compounds are represented on the top. **(b)** Representative micrographs on the left, depicting the results of the comet assay performed on yeast cells in the indicated conditions. Of note, NC and hydroxyurea (HU) represent the negative and positive controls, respectively. Violin plots on the right, depicting the comparative distribution of the comet tail length of ∼500 randomly selected cells per condition. Four different concentrations of 4NC and DP were used. Mann–Whitney U test was used to compute statistical significance between the test conditions and the negative control. **(c)** Schematic diagram on the left depicting the experimental workflow in a 96 well plate format for testing the mutagenic potential of the 4NC and DP. Of note, three different concentrations of 4NC and DP were used, and the mutant counts were normalized using Optical Density (O.D.) values at 600 nm. Representative micrographs in the middle, depicting the mutagenic effect of 4NC and DP in conferring canavanine resistance mutations, identified using spot assay (# of biological replicate = 52). Bean plot on the right depicting the distribution and mean of the normalized frequency counts of the canavanine resistant mutants, represented as fold change with respect to NC in the indicated conditions. The pairwise one-sample Student’s t-test was used to compute statistical significance between the test conditions and the negative control. **(d)** Schematic representation of the experimental design used in the quantitative estimation of reactive oxygen species (ROS) using the DCFH-DA dye-based assay. Box plot at the bottom depicting the ROS levels measured using DCFH-DA dye-based assay in the indicated conditions. Mann–Whitney U test was used to compute statistical significance between the test conditions and the negative control (NC). Notably, hydrogen peroxide (H_2_O_2_) treated yeast cells were used as a positive control. **(e)** Schematic representation of the experimental design of the high-throughput mutagenesis rescue experiment. Of note, 10 µM concentration of 4NC was used, with 50 µM L-ascorbic acid as ROS scavenger. Representative micrographs in the middle depicting the rescue of mutagenic effects of 4NC in the presence of ROS scavenger (# of biological replicate = 56). Bean plot on the right side depicting the distribution and mean of the normalized frequency counts of the canavanine resistant mutants, represented as fold change with respect to NC in the indicated conditions. The pairwise one-sample and two-sample Student’s t-tests were used to compute statistical significance between NC vs 4NC and 4NC vs 4NC+ROS scavenger, respectively.

To further evaluate the other predictions of Metabokiller, we next tested both DP and 4NC for inducing an anti-apoptotic response. For the apoptosis assay, we used a widely adopted, yeast-based assay, in which both the wild-type yeast cells and *Δfis1* knockouts were treated with the metabolites for 24 hours. We used acetic acid (AA) treatment as a positive control since it is known to induce apoptosis or programmed cell death in yeast. We observed induction of programmed cell death in the wild-type yeast cell treated with DP, 4NC, and acetic acid, whereas comparatively fewer cell deaths were observed for DP/4NC-treated *Δfis1* knockout cells than acetic acid, suggesting the activation of anti-apoptotic response by these metabolites, but not by acetic acid (**Figure 4a**). Notably, these results are largely in line with the Metabokiller predictions, further advocating the robustness of Metabokiller’s interpretability feature. Collectively, these results highlight the utility of Metabokiller in deciphering the putative mechanism of carcinogenicity induction by 4NC by virtue of its interpretability module.

**Figure 4:**
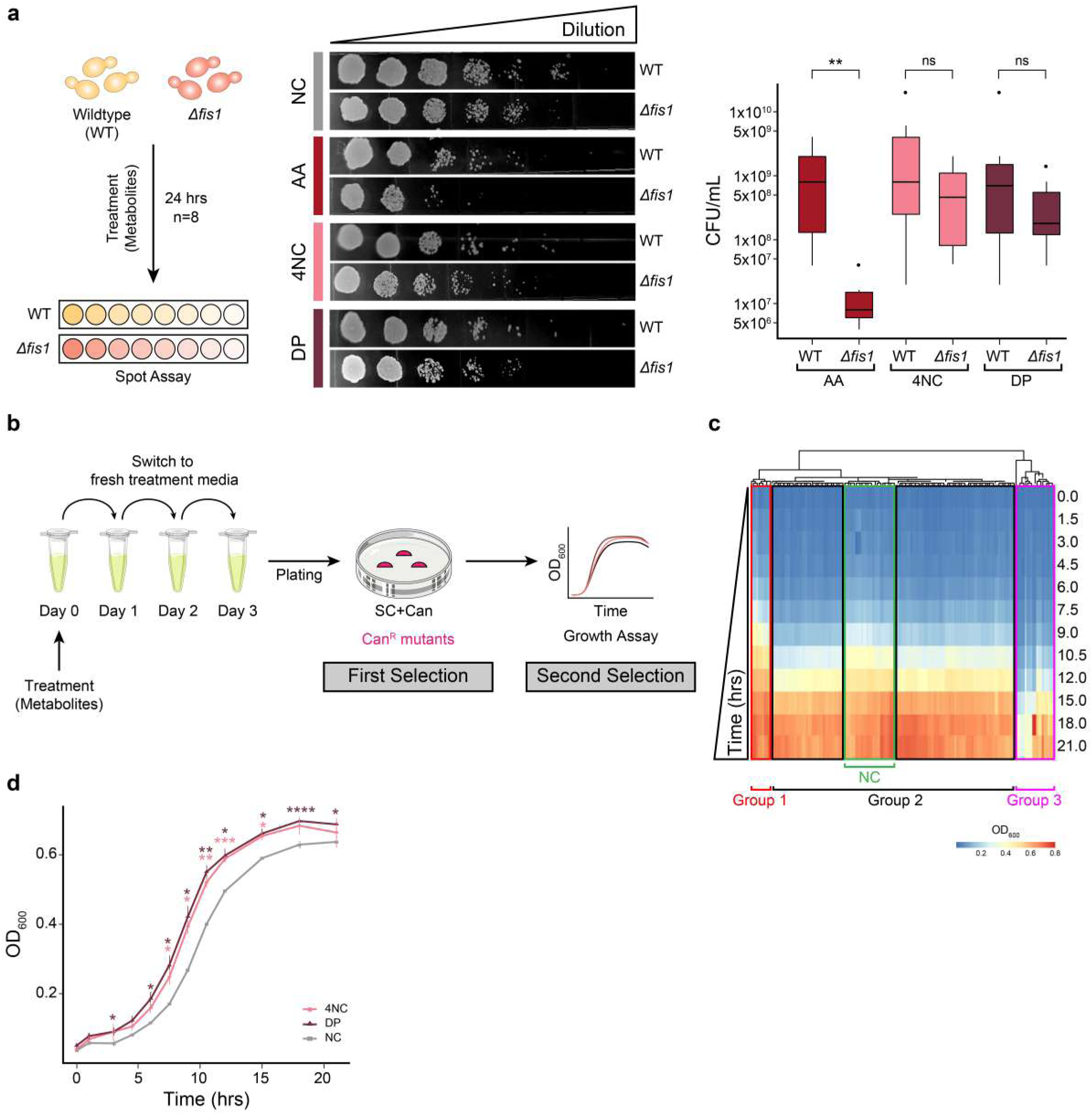
4-Nitrocatechol (4NC) and 3,4-Dihydroxyphenylacetic acid (DP) trigger an anti-apoptotic response in yeast. **(a)** Schematic representation of the experimental design of the apoptosis assay. Of note, 10 µM concentration of 4NC and DP was used. Representative micrographs in the middle depicting the DP and 4NC induced anti-apoptotic response in yeast. Untreated cells were taken as negative control (NC), while cells treated with 199 mM acetic acid (AA) for 200 min were used as a positive control. Box plot on the right highlighting the distribution of colony-forming units per mL after treatment of wild type (WT) and *Δfis1* yeast with vehicle (NC), acetic acid (AA), 3,4-Dihydroxyphenylacetic acid (DP), and 4-Nitrocatechol (4NC). Mann Whitney U test was used to compute the statistical significance with a p-value cutoff of < 0.05. **(b)** Diagrammatic representation of the experimental design used for the selection of gain of function mutants possessing accelerated cell division. Of note, mutants were selected based on canavanine resistance (selection 1) as well as accelerated cell division (selection 2). **(c)** Heatmap depicting the growth profiles of the metabolite-induced canavanine-resistant mutants. Mutants were clustered based on their growth profiles. Three major clusters were obtained with either accelerated (Group 1), decelerated (Group 3), or normal growth (Group 2). The green-colored rectangle in the middle indicates the growth profiles of the untreated cells (NC). **(d)** Growth curve profiles of untreated wild-type (NC), 4NC, and DP treated mutants with accelerated growth under optimal growth conditions. The Student’s t-test was used to compute statistical significance between the test conditions and the negative control (NC) with a p-value cutoff of < 0.05.

Since Metabokiller predicted both of these human metabolites as potential carcinogens, we investigated whether the mutations caused by 4NC and DP can alter the genes associated with cell cycle regulation, DNA repair response pathways, or other pathways related to carcinogenesis. We performed two consecutive selection procedures. In the first selection, we selected canavanine resistant mutants which in turn confirms the mutagenic effect of the tested metabolites. These mutants (n=167) were subjected to the second selection to assess alterations in the growth assay (**Figure 4b**). After obtaining the growth profiles of these mutants as well as a few randomly selected wild-type clones (omitted for selection 1), we performed hierarchical clustering based on their growth kinetics. Interestingly, we observed three clusters with yeast colonies displaying (a) accelerated growth (Group 1) (b) similar growth patterns as that of wild-type yeast (Group 2), and (c) decelerated growth (Group 3) **(Figure 4c, d)**. These results indicate that both 4NC and DP cause random mutagenesis and therefore result in distinct phenotypes (growth kinetics). In summary, all the aforementioned results collectively advocate for the accurate prediction by Metabokiller.

### Metabokiller predicted carcinogenic metabolites undergo mutation of cancer-associated genes and trigger malignancy

To investigate the impaired molecular mechanisms triggered by DP and 4NC treatment, we performed paired-end deep RNA sequencing on wild-type and mutant yeast cells in biological triplicates. Notably, only those mutants were selected that were canavanine-resistant (first selection) and displayed accelerated growth kinetics (second selection), reminiscent of cancer cells (**Figure 5a**). Differential gene expression analysis **(Figure 5b, c, Supplementary Figure 5 a-f)** on the normalized read counts identified a large number of downregulated genes, both in the case of DP and 4NC with respect to wild-type controls (**Figure 5d**). No overlap was observed in the upregulated genes, however, 35 downregulated genes were common between the conditions, suggesting differences in their mode of action at the molecular level (**Figure 5e**). We next attempted to decipher the impacted functional pathways impaired in these mutants using Gene Ontology Slim Term Mapper. Functional analysis of impaired genes revealed their involvement in DNA repair (*RAD59* and *DDR48*), chromatin organization (*ESC8 and RLF2*), DNA recombination (*NDJ1*, *RAD59*, and *DDR48*), and cell cycle (*YPR015C*, and *HUG1*). In addition to these, genes involved in cellular stress response were altered in both cases **(Figure 5f, Supplementary Figure 5g, Supplementary Table 8)**. Recent reports showed the possibility of inferring insertion-deletion (indel) mutations using RNA sequencing^73^. We next tracked indel mutations in wild-type and mutants using the raw FASTQ files. Mutational analysis identified specific indel mutations in the *CAN1* gene, validating the first selection procedure **(Supplementary Figure 5h)**. Using conservative filtering criteria (supporting read counts ≥ 5), we identified mutations in multiple protein-coding genes that are known to be involved in regulating DNA repair pathways as well as the cell cycle (**Figure 5g, h**). These independent analyses further support and explain the phenotypic response of the mutants (both 4NC and DP). Finally, we tested the carcinogenic effects of DP and 4NC on BEAS-2B normal human epithelial cells using soft-agar assay, a hallmark functional test for detecting malignant transformation. Our results clearly demonstrate that both 4NC and DP can induce malignant transformation in these cells, which collectively reinforce their carcinogenic potential (**Figure 5i-k**). Taken together, a combination of transcriptome and mutational analysis, coupled with a malignant transformation assay strongly suggests the carcinogenic role of 4NC and DP via genotoxicity.

**Figure 5:**
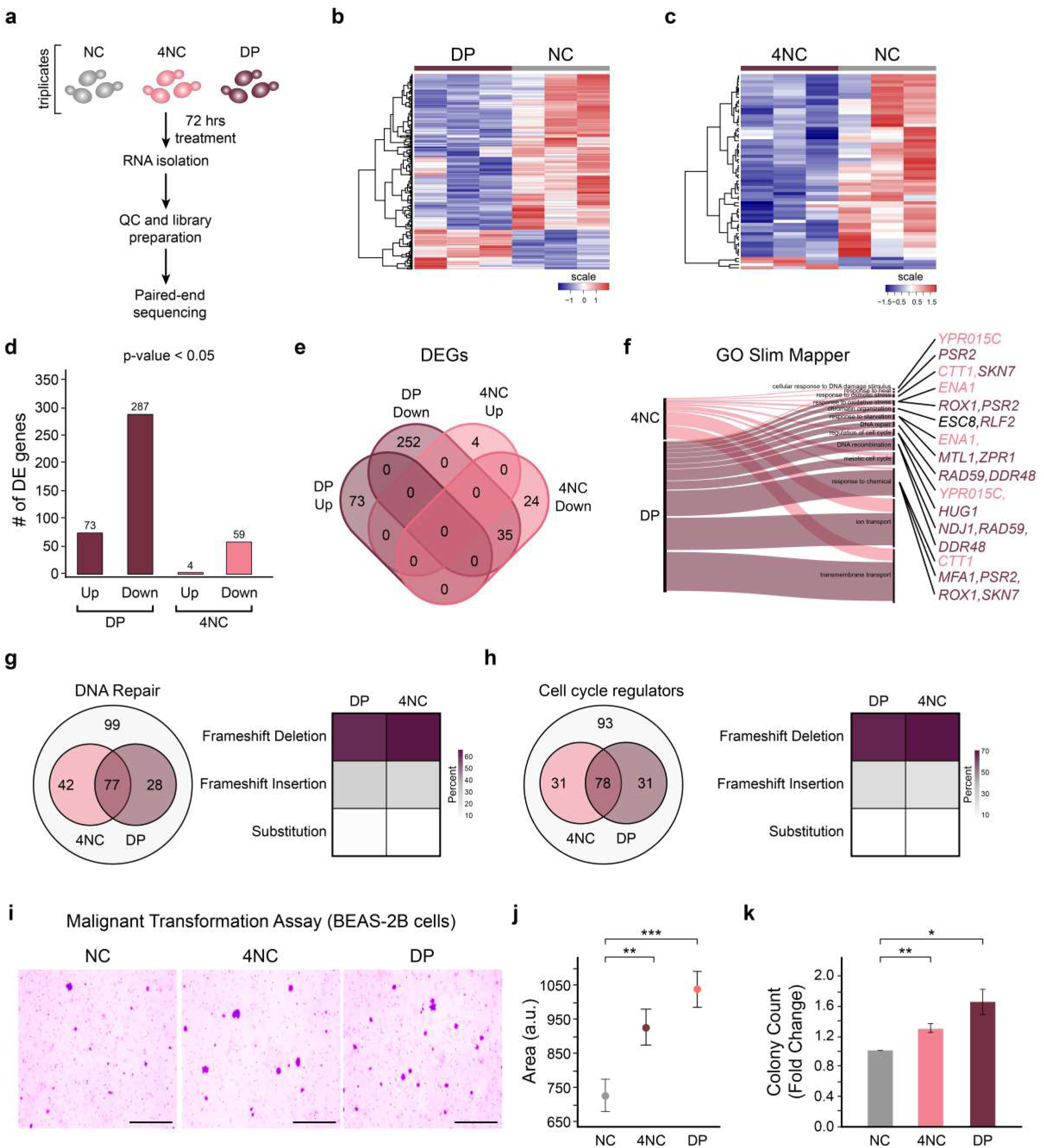
Cross-validation of carcinogenic and mutagenic potential of 4-Nitrocatechol (4NC) and 3,4-Dihydroxyphenylacetic acid (DP) using Deep RNA Sequencing and Soft Agar Assay. (**a**) Schematic representation depicting the experimental design of RNA sequencing, featuring the group information, treatment duration, and the sequencing parameters. (**b**) Heatmap representing the expression profile of differentially expressed genes in 3,4-Dihydroxyphenylacetic acid (DP) and untreated conditions (NC). **(c)** Heatmap representing the expression profile of differentially expressed genes in 4-Nitrocatechol (4NC) and untreated conditions (NC). **(d)** Bar graph depicting the number of differentially expressed genes in the indicated conditions. **(e)** Venn diagram depicting the number of differentially expressed genes shared among 4NC and DP (compared with respect to NC). **(f)** Gene Ontology Slim Mapper analysis depicting the association of the differentially expressed genes in the indicated ontologies. **(g-h)** The Venn diagrams and their accompanying heatmaps on the right side collectively depicting the number of mutated genes (insertions, deletions, and substitution mutations) involved in DNA repair or Cell Cycle regulation. The heatmap on the right further segregates the identified mutations as frameshift (insertion or deletion) and substitution. **(i)** Representative micrographs depicting the results of malignant transformation assay. Scale bar = 100 μm. **(j)** Mean-Whisker plot depicting the size of the malignantly transformed colonies in the indicated conditions. Two sample Student’s t-test was used to compute the statistical significance between test conditions and the negative control (NC), with a p-value cutoff of < 0.05. **(k)** Bar graphs represent the fold change increase in the number of transformed colonies in the indicated conditions. Two sample Student’s t-test was used to compute the statistical significance between test conditions and the negative control (NC), with a p-value cutoff of < 0.05.

## DISCUSSION

The integrity of a cellular genome is always vulnerable to both intrinsic and extrinsic factors^74,75^. A large degree of effort is ongoing to delineate the compositional chemistries of the exogenous compounds that possess a direct threat to genomic fidelity^74^. However, recent evidence suggests an equal degree of threat from the endogenous metabolites^76,77^. Irrespective of the source, the induced DNA alterations have been implicated in cancer, as well as aging^78,79^. Therefore, accurate evaluation of the carcinogenic potential of any compound is immensely important and plays a vital role in decreasing the cancer burden. Despite the availability of a stringent mandate for carcinogenicity testing, many of the Food and Drug Administration (FDA) approved drugs have been identified as carcinogens and therefore later withdrawn from the market^80^. As a precautionary measure, stringent protocols have been set up to test the proposed drugs for their carcinogenicity potential. The QSAR models have been used for the computation-assisted pre-screening for carcinogenicity^21,23,24,31,32^. Though multiple computational approaches have been used for the carcinogenicity predictions, unfortunately, they largely fall short on unseen data, primarily due to the limited amount of experimentally-validated data available for model training. In this work, we have used a novel approach where, unlike utilizing limited information on *bona fide* carcinogens and non-carcinogens, we leverage biochemical properties associated with known carcinogens. Our method, Metabokiller, outperformed most of the recent and widely used methods for carcinogenicity prediction. Moreover, in addition to the higher prediction accuracy, Metabokiller also possesses features of explainable artificial intelligence, since it provides the individual contribution of all the six-core models detailing the biochemical properties of carcinogenicity. We used Metabokiller to perform a large-scale computational screening of human metabolites for their carcinogenic potential. We identified a large number of previously known and unknown human metabolites with carcinogenicity potential. Our functional analysis of carcinogen-predicted metabolites on pathway space revealed selective enrichment of tyrosine and tryptophan metabolism, nucleotide metabolism, metabolic pathways associated with drug metabolism, xenobiotics, oxidative stress, etc. Further, functional over-representation analysis of predicted carcinogenic metabolites identified 38 previously known carcinogenic metabolites. These results strongly advocate for the robustness of biochemical features in capturing carcinogenicity. Our in-depth investigation of tumor-associated metabolites leads to the discovery of a large number of previously uncharacterized oncometabolites. Of note, in addition to these, Metabokiller also flagged well-characterized oncometabolites such as 2-hydroxyglutarate^69^, succinate, fumarate^81,82^, adenylosuccinate, and polyamines (putrescine, spermidine, and spermine)^66^. Fumarate is a metabolic intermediate of the Krebs cycle. Functional studies involving loss of function of fumarate hydratase (FH) enzyme suggest that the accumulation of fumarate triggers renal cell cancer^81,82^. Moreover, similar to fumarate, both succinate and adenylosuccinate are also linked to the same pathway. Loss of function studies involving succinate dehydrogenase (SDH), a critical enzyme that regulates cellular succinate levels suggest that accumulation of succinate triggers cancer, and tumor repopulation post radio/chemotherapy^81–83^. Accumulation of 2-hydroxyglutarate rewires the cancer cell metabolism and triggers oxidative stress. In addition to its direct interference with the metabolism, it also acts as a modulator of chromatin remodeling enzymes (2-oxoglutarate-dependent dioxygenases), hypoxia-inducible factor (HIF), and mammalian target of rapamycin (mTOR) pathways leading to carcinogenesis^69^. While the functional characterization of oncometabolites provides an orthogonal dimension to uncovering tumor biology, to date, only a handful of metabolites have been characterized as oncometabolites. Metabokiller offers a unique approach to uncovering oncometabolites and also provides a biochemical property-driven explanation for their mode of action.

Despite possessing multiple advantages and immunity towards limited training data for carcinogenicity prediction, Metabokiller also possesses several limitations. First, it has been known that carcinogens also possess other biological properties such as induction of chronic inflammation, inhibition of senescence, cell transformation, changes in growth factors, energetics, signaling pathways related to cellular replication, or cell cycle control, angiogenesis, and are immunosuppressive in nature^84^. The present ensemble model does not take into account these properties. This is mainly due to the paucity of these compounds in the available literature. Second, an inert compound can also harbor carcinogenic properties after being processed by the cellular enzymes^43^. These assumptions are also not taken into consideration in the present ensemble model of Metabokiller. Third, the carcinogenicity of certain molecules is dose-dependent, however, Metabokiller does not factor in the dosage information for predictions. Fourth, unlike other proposed toxicogenomic-based prediction models, Metabokiller does not account for tissue specificity and merely relies on the carcinogenicity status^85,86^. Lastly, the present version of Metabokiller only supports carcinogenicity prediction and does not allow assessment of the toxicity properties such as oral toxicity, liver toxicity, etc. Irrespective of these aforementioned limitations, Metabokiller significantly satisfies the urgent need for a robust, reliable, and accurate alternative for carcinogenicity prediction. Furthermore, the interpretability module of Metabokiller provides a biochemical enriched explanation for each prediction (**Figure 6**). The high Recall value of Metabokiller, an essential feature for any health-related prediction model, allows accurate identification of carcinogens (true positives). Large-scale screening of the human metabolome by Metabokiller provides high confidence predicted carcinogenic metabolites and opens a new avenue in functional metabolomics and may contribute to unfolding the role of cancer-associated metabolites in disease progression.

**Figure 6:**
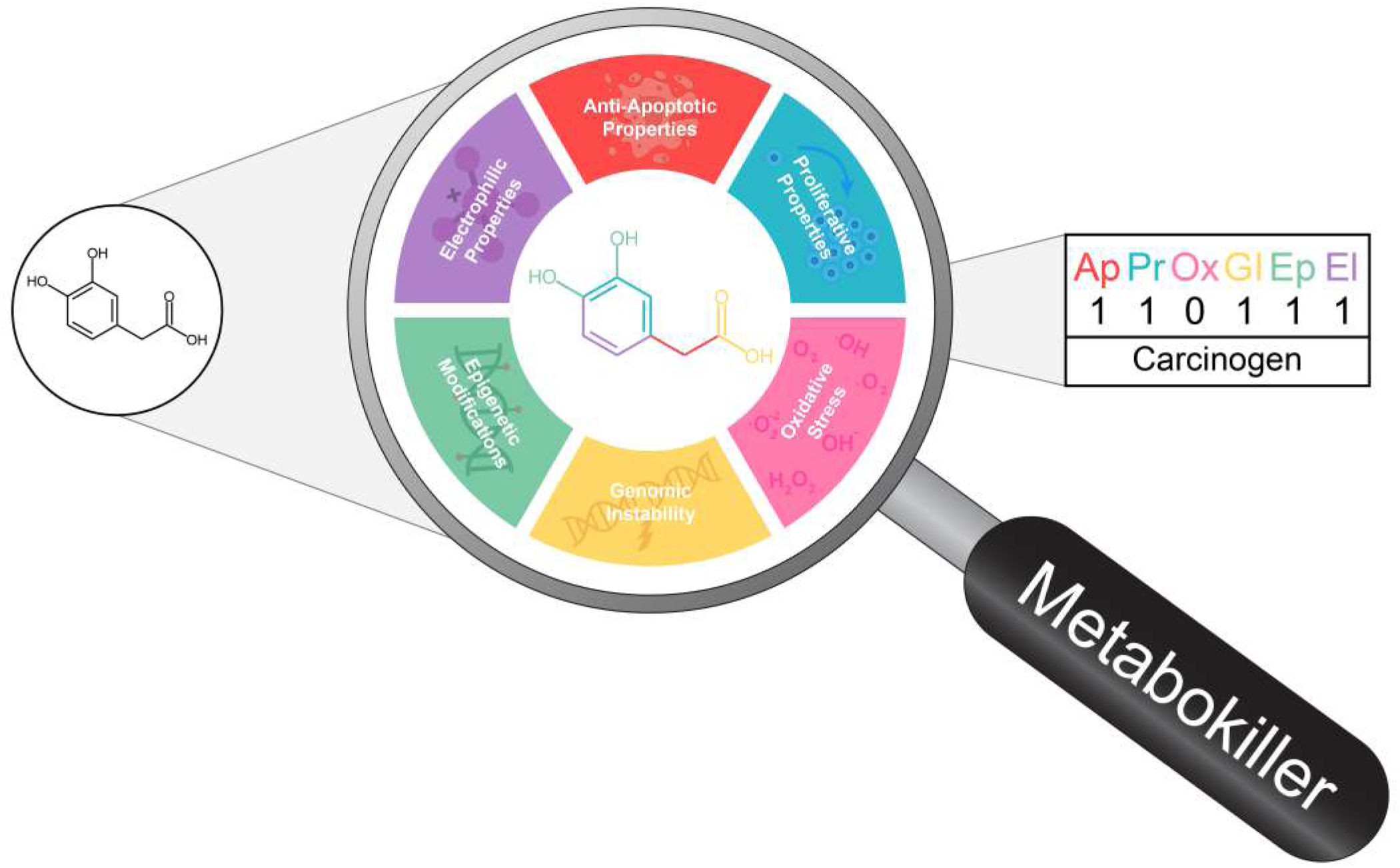
Graphical Abstract of the key findings. Schematic representation depicting the mode of action of Metabokiller. Of note, Metabokiller predicts the carcinogenicity potential of any query compound by assessing for the indicated biochemical properties.

## MATERIALS AND METHODS

### Data compilation

To build an Ensemble model (Metabokiller) for carcinogenicity prediction, we opted to utilize both the biological and chemical properties associated with carcinogens. These include the potential to induce a proliferative response, cellular oxidative stress, epigenetic alterations, genomic instability, and anti-apoptotic properties. Since carcinogens attack DNA by virtue of their electrophilic nature^12,44^, we, therefore, selected electrophilic potential as one of the properties. The training data for the six independent models were compiled from the literature or numerous databases in three consecutive steps (combined training data is annotated as MK_Tn_). Of note, data curation, filtering, and cross-validation were achieved manually to ensure its authenticity. In the first phase, we compiled data from all the available literature, databases, and web servers. In the second phase, we assimilated both the pro- and anti- or no-activity compounds for each of the six models (**Supplementary Table 1**). Finally, in the third phase, we filtered, cross-validated, thoroughly rechecked, and removed the contradictory entries. Post these stringent screening steps, we gathered a total of 12957 positive (Class 1) and 22711 negative (Class 0) compounds. In summary, we obtained 12127 compounds for genomic instability model (class 0 = 8931 and class 1 = 3196), 1714 compounds for electrophilic property model (class 0 = 746 and class 1 = 968), 15145 compounds for epigenetic modification model (class 0 = 7576 and class 1 = 7569), 798 compounds for anti-apoptotic property model (class 0 = 525 and class 1 = 273), 225 compounds for proliferative property model (class 0 = 123 and class 1 = 102) and 5659 compounds for oxidative stress model (class 0 = 4810 and class 1 = 849). A detailed description of all compounds in the respective datasets is mentioned in Supplementary Table 1. In addition to the aforementioned datasets, we also gathered a list of experimentally-validated carcinogens and non-carcinogens from multiple sources and subsequently annotated them as Metabokiller External Training data (MK_ETn_), Metabokiller Testing data 1 (MK_Ts1_), Metabokiller Testing data 2 (MK_Ts2_), and Independent Dataset (I.D.). Detailed information about the compounds in these datasets is mentioned in Supplementary Table 2. Simplified molecular-input line-entry system (SMILES) were extracted for all the molecules using PubChem Identifier Exchange Service (https://pubchem.ncbi.nlm.nih.gov/idexchange/idexchange-help.html), which were further converted into canonical SMILES using OpenBabel^87^.

### Model Building

The core of the Metabokiller ensemble model relies on the six independent predictive models that collectively define the biochemical space of carcinogens. Each model uses independent training data that is further segregated into two classes based on their known activity i.e. class 0: anti- or no-activity and class: 1 pro-activity. For all the models, the training dataset was evaluated for class imbalance, and subsequently, the upsampling/downsampling techniques were used to counteract this bias. We initially tested three different feature extraction methods i.e. bioactivity-based descriptors (Signaturizer library), chemistry-based molecular descriptors (Mordred software), and Graph-based (DeepChem library). We also tested multiple classification algorithms i.e., Random Forest (RF), Multilayer perceptron (MLP), k-Nearest Neighbor (KNN), Support Vector Machine (SVM), Stochastic Gradient Descent (SGD), Logistic Regression (LR), GraphConvModel (GCM), Attentive FP, Graph Convolution Network (GCN), and Graph Attention Network (GAT) for model building. Notably, for feature selection, we used Boruta, a feature selection algorithm, and for down/upsampling, Synthetic Minority Oversampling Technique (SMOTE) was used^88^. Briefly, we tried all the aforementioned combinations and built twelve distinct models for each of the biochemical properties using the default parameters. We next selected the best performing models for each of the biochemical properties and subsequently performed the random-grid-search hyperparameter tuning to obtain the best and most stable hyperparameters for model building. We finally trained the models with the best hyperparameters and rigorously evaluated their performance by using both the bootstrapping (20 folds) as well as 10-fold cross-validation techniques. A dataset comprising a list of *bona fide* experimentally-validated carcinogens/non-carcinogens (MK_ETn_) was used to build the ensemble model. For this, we projected the MK_ETn_ dataset on all the six biochemical models and obtained their prediction probabilities for pro-activity. We next used these probability values as features to build the ensemble model. Of note, we have selected Gradient Boosting Machine-based model since it outperformed other models in prediction performance. We further improved its performance by performing random-grid-hyperparameter tuning **(Supplementary Table 4)**. Finally, to negate the impact of data bias, we selected twenty distinct GBM-based models generated using Bootstrapping and implemented a majority voting-based method for carcinogenicity prediction. To provide quantitative means for model interpretability, we have also implemented Local Interpretable Model-agnostic Explanations (LIME).

### Strains

The BY4741 strain (MATa *his3Δ1 leu2Δ0 met15Δ0 ura3Δ0*) of *Saccharomyces cerevisiae* was used in all the experiments. For the Apoptosis assay, *Δfis1* (MATa *fis1::*kanMX4 *Δhis3Δ1 leu2Δ0 met15Δ0 ura3Δ0*) knockout yeast strains were used. Unless mentioned, the yeast is grown at 30°C at 200 Revolutions Per Minute (rpm) in the Yeast Extract–Peptone–Dextrose (YPD) (1% Yeast extract, 2% Peptone, 2% Dextrose). 1.5% of Agar is additionally added to YPD to prepare plates.

### Comet Assay

Yeast secondary culture with an estimated optical density of 0.6-0.7 (OD_600_) was used for the comet assay. Briefly, yeast in the secondary culture was exposed to different concentrations of 4-Nitrocatechol (4NC) (N15553, Sigma-Aldrich), and 3,4-Dihydroxyphenylacetic acid (DP) (11569, Sigma-Aldrich) at the final concentrations of 0.1 μM, 1 μM, 10 μM, and 100 μM. Moreover, hydroxyurea and solvent alone were used as positive and negative controls, respectively. The yeast cells were grown in the presence of metabolite/compounds for 90 minutes at 30°C at 200 rpm. Post-treatment, an equal number of cells were taken from each condition and pelleted down using centrifugation. Pellets were washed twice with pre-cooled 1X Phosphate-buffered saline (PBS) to remove media traces. The pellet was resuspended in 0.3% w/v agarose prepared in S-Buffer (1 M sorbitol, 25 mM KH2PO4, pH 6.5). Next 40 units of zymolyase enzyme (L2524, Sigma-Aldrich) was added to degrade the cell wall. Finally, 80 μL of this solution was spread onto glass slides pre-coated with 0.8% w/v agarose. Slides were incubated at 30°C for 30 min for cell wall degradation. Finally, the slides were incubated in the freshly prepared lysis solution (30 mM NaOH, 1 M NaCl, 0.05% w/v SDS, 50 mM EDTA, 10 mM Tris–HCl, pH 10) for 2 hours. The slides were washed three times for 20 min each with electrophoresis buffer (30 mM NaOH, 10 mM EDTA, 10 mM Tris–HCl, pH 10) to remove the lysis solution. Next, an electrophoresis step was performed in the electrophoresis buffer for 15 min at 70 mV/cm. Post electrophoresis, the slides were incubated in the neutralization buffer (10 mM Tris–HCl, pH 7.4) for 10 mins, followed by 10 minutes of incubation in 76% and then 96% ethanol. After this, Giemsa stain was added for 20 min to stain the DNA. The slides were then washed thrice with 1x PBST (0.1% Tween20) for 10 min each. The comets were visualized in a brightfield microscope. Micrographs were taken from random sections of the slides. Approximately 500 cells were chosen randomly from each condition and the comet tail length was measured using ImageJ software (https://imagej.nih.gov/).

### Mutagenesis Assay

Yeast was grown overnight at 30°C at 200 rpm in YPD media. An equal number of cells were inoculated (measured using OD_600_) into 96 well plates containing 100 μL of YPD media along with respective metabolites (4NC and DP) at concentrations of 0.1 μM, 1 μM, and 10 μM each. The solvent alone was used as a negative control. A total of 52 biological replicates were set up for each condition. The 96 well plates were covered with a Breathe-Easy^®^ sealing membrane (Z380059-1PAK, Sigma-Aldrich) to prevent media evaporation. Plates were incubated at 30°C at 200 rpm and the media was replenished with fresh media 24 hours post-incubation with the aforementioned metabolites or vehicles. Post 48 hours of incubation, cell count was estimated using optical density (OD_600_) measurements which were further used to normalize the Can^R^ mutant frequencies. Subsequently, a Spot assay was performed on Synthetic Complete Medium (2% dextrose, 6.7 g/L yeast nitrogen base without amino acids, 20 mg/L histidine, 120 mg/L leucine, 20 mg/L methionine, 20 mg/L uracil, 20 mg/L adenine, 2% agar) containing canavanine (60 μg/mL) agar plates. The plates were incubated at 30°C for 2-3 days, the number of colonies were counted, and subsequently normalized using the OD_600_ measurement values from the aforementioned steps. The normalized mutant frequencies were plotted using R Programming. A similar setup was used for the rescue experiments, where the cells were grown along with 4NC at a final concentration of 10 μM, in the presence and absence of a ROS scavenger, i.e. 50 μM L-Ascorbic acid.

### Cell Death Assay

Propidium Iodide staining was used for the quantitative estimation of the cell deaths in the treated and untreated conditions. Cells were grown overnight at 30°C at 200 rpm in YPD media. Using this primary culture as inoculum that ensures equal cell counts, cells were grown in 96 well deep plates in the presence of respective metabolites (4NC and DP) at concentrations of 0.1 μM, 1 μM, and 10 μM each for 12 hours with 8 biological replicates. Propidium iodide staining was performed on aliquots after nine and twelve hours of incubation in 96 well fluorescence plates. Heat-killed cells were used as a positive control. Propidium iodide (11195, SRL) was added at a final concentration of 5 μg/mL and incubated in the dark for 15 minutes. The fluorescence was measured using Biotek Synergy HTX multi-mode reader at excitation and emission wavelengths of 530/25 nm and 590/25 nm, respectively.

### Apoptosis Assay

Wild type BY4741 and *Δfis1* (MATa *fis1::*kanMX4 *Δhis3Δ1 leu2Δ0 met15Δ0 ura3Δ0*) knockout yeast strains were grown overnight at 30°C at 200 rpm in YPD media. An equal number of both wild-type and *Δfis1* cells (optimized using OD_600_) were inoculated into multiple 96 well plates containing 100 μL of YPD media along with respective metabolites (4NC and DP) at a concentration of 10 μM for 24 hours with 8 biological replicates. Cells treated with 199 mM acetic acid for 200 min were used as a positive control (AA), and untreated cells as negative control (NC). Post-incubation, serial dilutions of 1:10 were prepared in distilled water, and a Spot assay was performed on Synthetic Complete Medium agar plates. The plates were incubated at 30°C for 2-3 days, the number of colonies werevcounted, and subsequently, Colony Forming Units per mL of media (CFU/mL) were calculated using the formula^89^:

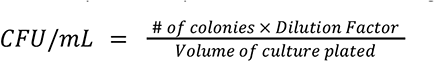

### Reactive Oxygen Species Quantification Assay

2’-7’dichlorofluorescein diacetate (DCFH-DA) staining was used to measure the levels of intracellular reactive oxygen species. Yeast was grown overnight at 30°C at 200 rpm in YPD media. Using this primary culture as inoculum that ensures equal cell counts, cells were grown in 96 well deep plates in the presence of respective metabolites (4NC and DP) at concentrations of 0.1 μM, 1 μM, and 10 μM each for 12 hours with 8 biological replicates. DCFH-DA fluorescence staining was performed after nine and twelve hours of incubation in a 96-well fluorescence plate with 10 mM hydrogen peroxide as a positive control. DCFH-DA (85048, SRL) was added at a final concentration of 10 μM in each well. The plate was incubated in the dark for 30 minutes. The fluorescence was measured using Biotek Synergy HTX multi-mode at excitation and emission wavelengths of 485/20 nm and 525/20 nm, respectively.

### Selection of Canavanine-resistant mutants with Accelerated Cell Division

Yeast (BY4741) was grown at 30°C at 200 rpm in YPD media. Using this primary culture as inoculum, cells were grown in MCTs in the presence of respective metabolites (4NC and DP) at concentrations of 0.1 μM, 1 μM, and 10 μM each for 72 hours with 4 biological replicates, alongside negative control. The culture was centrifuged and resuspended in fresh media after every 24 hours. After 72 hours, the cells were plated onto Synthetic Complete Medium containing canavanine (30 μg/mL) agar plates to screen for canavanine-resistant colonies. This is referred to as the first round of selection. The canavanine-resistant colonies from all the conditions were grown in YPD media in the presence of canavanine but without metabolites, and a second selection was performed using the growth assay. RNA was isolated from the canavanine resistant mutants harboring accelerated growth, and post quality check was used for library preparation and sequencing.

### RNA isolation

Cells (BY4741) were grown at 30°C at 200 rpm in Synthetic Complete Medium containing canavanine (30 μg/mL) in biological triplicates and harvested and resuspended in 500 μL of 1x PBS. Zymolyase (40 units) treatment was performed by incubating for 30 min at 30°C. Post zymolyase treatment, cells were lysed by mechanical breakdown using the glass beads. The lysed cell suspension was transferred to fresh MCT, centrifuged and the pellet was resuspended in 500 μL of TRIzol™ reagent (15596026, Ambion life technologies). 500 μL of chilled chloroform was added to each tube, followed by 15 min of mechanical breakdown and 5 minutes of incubation at room temperature. Samples containing TRIzol™ and chloroform were centrifuged at 12000 g for 15 minutes at 4°C. The upper aqueous layer was carefully transferred to a fresh MCT containing 250 μL of prechilled isopropanol. 1 μL of glycogen was also added to accelerate RNA precipitation, followed by transient incubation for 30 minutes at -80°C. Nucleic acid was pelleted down by centrifugation at 12000 g for 15 min at 4°C. Pellet was washed with 70% ethanol at 5000 g for 5 minutes at 4°C and air dried, and subsequently resuspended in RNase-free water. To remove DNA contamination, samples were treated with DNase at 37°C for 15 minutes. DNase inactivation was performed by heating the samples at 65°C with 5 mM EDTA for 10 minutes. This purified RNA was resuspended in RNase-free water for downstream processing.

### RNA Sequencing Analysis

Post sequencing the quality control check was performed using MultiFastQ. The paired-end sequencing read files were mapped to the yeast reference genome (ENSEMBL; R64-1-1; GCA_000146045.2) using the align function of the Rsubread package (v2.6.4)^73^, and the aligned BAM files were generated. Trimming of reads was performed from both ends prior to mapping. We used the inbuilt function of Rsubread to detect short indel (Insertions and Deletions) mutations and the resulting information was stored in the Variant Call Format (VCF) format. An expression matrix containing the raw read counts for each sample (**Supplementary Table 7**) was obtained using the featureCounts function. Read count matrix was normalized and analyzed using the RUVg function of the RUVSeq package (v1.24.0)^90^ to remove unwanted variation from RNA-Seq data. Differentially expressed genes were computed using the DESeq2 package (v1.30.1)^91^, with a fold change cutoff of 1.5, and a p-value significance of < 0.05. Functional Gene Ontology analysis was performed using the GO Slim Mapper tool of Saccharomyces Genome Database (SGD) (https://www.yeastgenome.org/goSlimMapper) and Metascape (https://metascape.org/gp/index.html).

### Mutation Analysis

Variant Call Format (VCF) files containing indel mutation information were generated using the Rsubread package as discussed above. We next merged the variant information from three replicates and a single VCF file was generated using the VCFtools (v0.1.16-1)^92^. For the downstream analysis, we only selected those mutations that carry the number of supporting reads for variants (SR) ≥ 5. Cross comparison of the VCF files from treatment groups of 4NC and DP was performed with that of negative control (untreated). Finally, we only selected those variants that were unique to each treatment condition. The mutations were segregated into frameshift (insertion or deletion) or substitution using the SIFT 4G Annotator using R64 (sacCer3) as a reference genome^93^.

### Soft Agar Assay

BEAS-2B cells were cultured in Dulbecco’s Modified Eagle Medium (DMEM) media supplemented with 10% FBS and 1% Penicillin-Streptomycin at 37°C humidified incubator with 5% CO_2_. For soft agar assay, BEAS-2B cell suspension of 1×10^4^ cells/well was added to the top agar solution at a final concentration of 0.3% and final volume of 0.5 mL per well and suspended on the top of solidified 0.5% agar base. The overnight culture was treated with 10 µM DP or 10 µM 4NC. The cells were supplemented with DMEM complete media twice until colonies appeared after 3 weeks. Colonies were stained with 0.1% crystal violet in 10% ethanol and imaged using a 4x lens and bright-field filter in the light microscope.

### Statistical Analysis

All statistical analyses were performed using either the Past 4 software or R-Programming. For comparison of the medians of the two distributions (non-parametric), Mann–Whitney U test was used, while two-sample Student’s t-test was used for comparing the means (pairwise comparisons). One sample Student’s t-test was used to measure the significance of the mutagenesis and its rescue experiments. The p-value cutoff used in this study is 0.05. *, **, ***, and **** refer to p-values <0.05, <0.01, <0.001, and <0.0001, respectively.

## SUPPLEMENTARY INFORMATION

**Supplementary Table 1:** Tabular representation of the datasets used for building the individual six models capturing the carcinogen-specific biochemical properties. Of note, each table contains information about the compound canonical SMILES, common name, PubChem CID, InChI, IUPAC name, activity status, and source information.

**Supplementary Table 2:** Tables containing the list of experimentally validated carcinogens and non-carcinogens that constitute training and testing datasets used in this study. Of note, each table contains information about the compound canonical SMILES, common name, PubChem CID, InChI, IUPAC name, carcinogenicity status, and source information.

**Supplementary Table 3:** Table containing performance metrics of all the tested classifiers for building the ensemble model (Metabokiller).

**Supplementary Table 4:** Table containing information about the hyperparameters used to build all the models supported by Metabokiller.

**Supplementary Table 5:** Table containing information about the carcinogenicity prediction results of the Human Metabolome Database (HMDB) metabolites using Metabokiller (probability cutoff ≥ 0.7). It also contains information about compound canonical SMILES, HMDB accession ID, PubChem CIDs, HMDB status, InChI keys, KEGG IDs, compound names, IUPAC names, and prediction probabilities of all the individual models supported by Metabokiller.

**Supplementary Table 6:** Table containing carcinogenicity or mutagenicity prediction results of 3,4-Dihydroxyphenylacetic acid (DP), 4-Nitrocatechol (4NC) using different predictive models/methods.

**Supplementary Table 7:** Table containing gene expression data (read counts) of canavanine resistant mutants treated for 72 hours with 3,4-Dihydroxyphenylacetic acid (DP), 4-Nitrocatechol (4NC), and vehicle (negative control).

**Supplementary Table 8:** Table containing information about the Gene Ontology results computed using the differentially expressed genes in the indicated conditions. The table also contains information about Gene Ontology IDs, Term names, and the information of the mapped gene for each of the given Ontology. Of note, this analysis was performed using the GO Slim Mapper tool from the Saccharomyces Genome Database (SGD) website.

### ETHICS APPROVAL AND CONSENT TO PARTICIPATE

Not applicable

## CONSENT FOR PUBLICATION

All authors have read and approved the manuscript for publication.

## DATA AVAILABILITY

The raw RNA sequencing files are available at ArrayExpress with Accession ID: E-MTAB-11179. The processed datasets are provided in the supplementary tables.

## CODE AVAILABILITY

A Python package for Metabokiller is provided via pip https://pypi.org/project/Metabokiller/ or from the project GitHub page: https://github.com/the-ahuja-lab/Metabokiller and Zenodo (10.5281/zenodo.6477021). Code used for building machine learning models is provided on the project GitHub page.

## COMPETING INTERESTS

A provisional patent has been filed (Reference No. 202111052929, Application No. TEMP/E-1/60118/2021-DEL) describing the computational architecture of the Metabokiller. Usage of the Metabokiller Python package is free for the academic institutions, or for any academic-related project, however, for commercial usage, users must contact the authors.

## FUNDING

The Ahuja lab is supported by the Ramalingaswami Re-entry Fellowship (BT/HRD/35/02/2006), a re-entry scheme of the Department of Biotechnology (DBT), Ministry of Science & Technology, Govt. of India, Start-Up Research Grant (SRG/2020/000232) from the SERB, Science, and Engineering Research Board and an intramural Start-up grant from Indraprastha Institute of Information Technology-Delhi (IIIT-Delhi). The Sengupta lab is funded by the INSPIRE faculty grant from the Department of Science & Technology (DST), India.

## AUTHORS’ CONTRIBUTIONS

The study was conceived by G.A. Computational analysis workflows were designed by G.A., D.S, and A.M. Yeast and Human experimental workflows were designed by G.A., A.M., and S.N., respectively. Yeast-based assays were performed by A.M., S.A., and N.D. Human cell culture-based experiments were performed by S.S. Data compilation for the model building was performed by A.M., P.G., A.A. P.R., and analysis workflow was made by S.M., V.G., S.A., A.M., R.S., R.G. and P.G. V.S., An.M. and J.T assisted in data interpretation. Metabokiller Python package was created by S.M. Illustrations were drafted by A.M. and G.A. G.A. and A.M. wrote the paper. All authors have read and approved the manuscript.

## ACKNOWLEDGEMENTS

The authors would like to thank the IT-HelpDesk team of IIIT-Delhi for providing assistance with the computational resources. We thank all the members of the Ahuja lab for their intellectual contributions at various stages of this project. We also thank Dr. Kaustuv Datta for providing critical insights into this study and Dr. Kausik Chakraborty for sharing yeast strains.

---

**Supplementary Figure 1:**
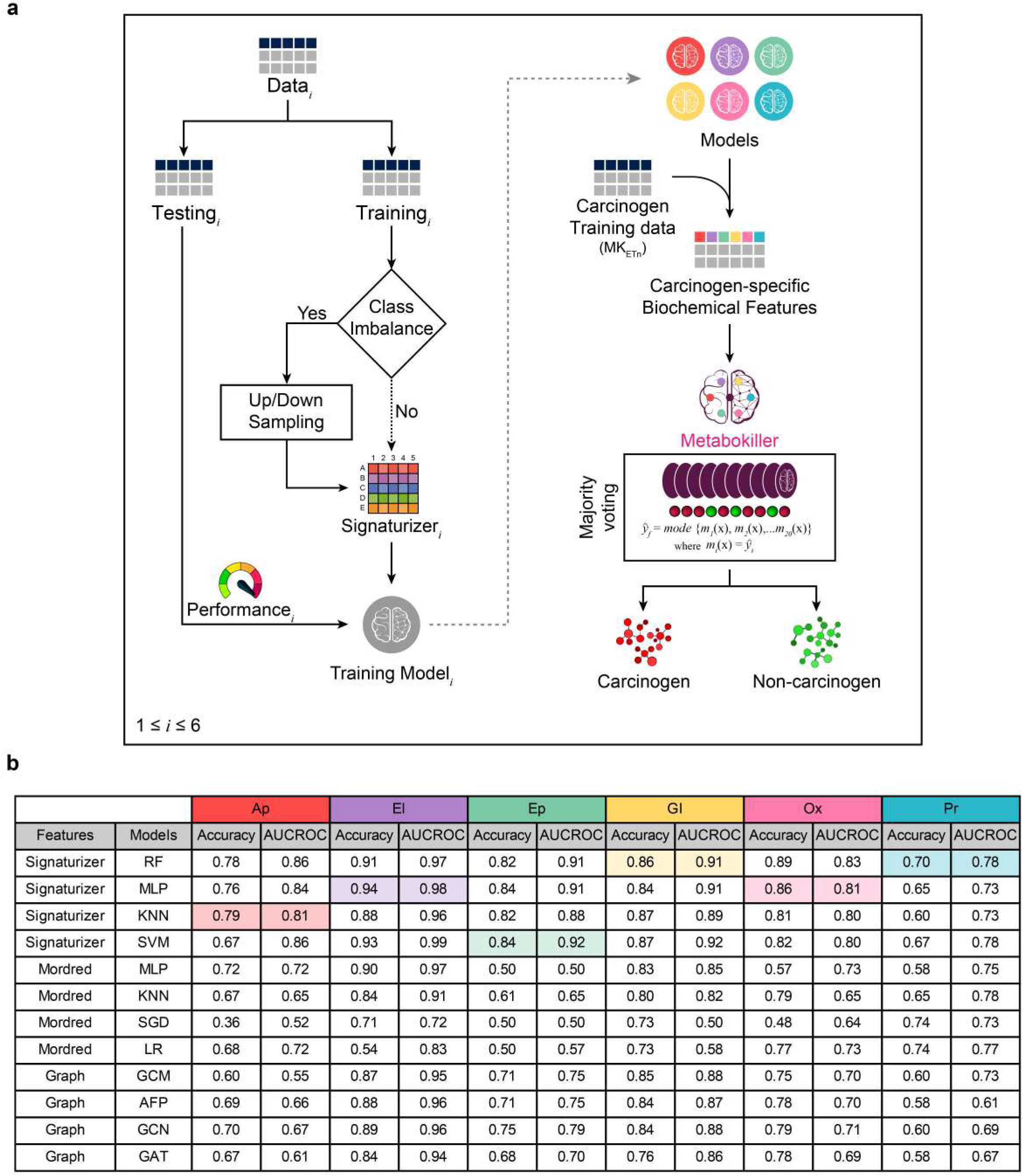
Metabokiller: An ensemble model that leverages biochemical properties to predict carcinogenicity. **(a)** Schematic representation depicting the step-by-step workflow used to build all the six individual biochemical models and the ensemble model (Metabokiller). Up/downsampling approach was used to counteract the class imbalance. Signaturizer library was used to generate bioactivity features. Hyperparameter tuning was performed to obtain the best-performing model parameters. The ensemble model (Metabokiller) was built using biochemical features of experimentally-validated carcinogens/non-carcinogens generated using six models. The majority voting method was used to assign the final carcinogenicity status. **(b)** The table containing performance metrics (Accuracy and AUCROC) of the twelve base models representing indicated biochemical properties. Of note, the models differ in their usage of feature extraction methods i.e. Signaturizer (bioactivity-based descriptors), Mordred (chemistry-based descriptors), and DeepChem library (graph-based descriptors) as well as classification algorithms. Models selected to build Metabokiller are color-highlighted.

**Supplementary Figure 2:**
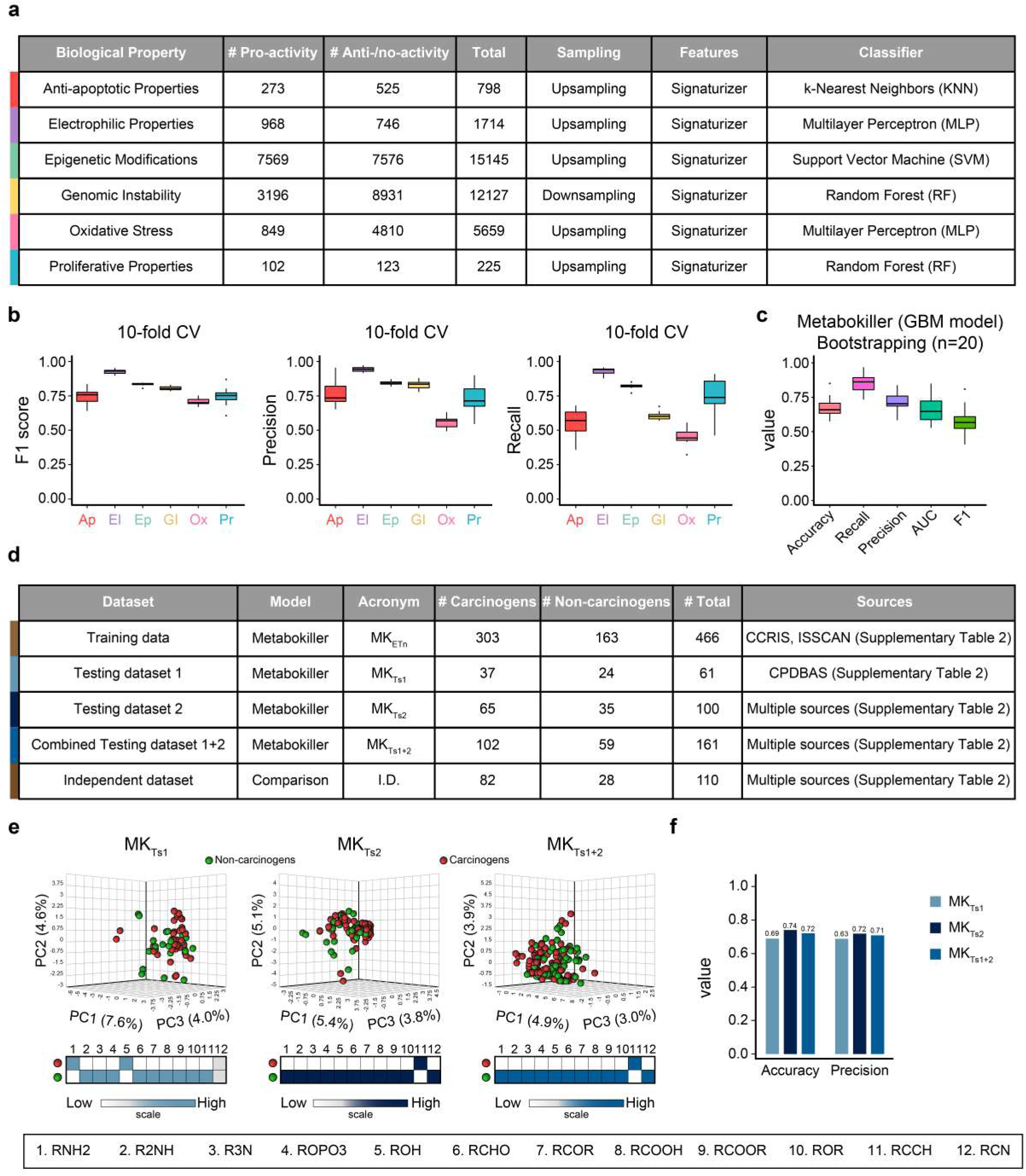
Metabokiller leverages multiple datasets to generate supporting models. (a) Table denoting the information about the number of compounds, the method used for handling class imbalance, the feature extraction method, and the classifying algorithm used to build the indicated models. (b) Box plots depicting the F1 Score, Precision, and Recall of the indicated models as inferred from the 10-fold cross-validation. (c) Box plot depicting the model performance of the twenty Gradient Boosting Machine (GBM)-based models generated using bootstrapping technique. (d) Table containing source and compound information about the datasets used in this study. (e) Principal Component Analysis revealing the chemical heterogeneity between the carcinogens and non-carcinogens in the indicated datasets. The heatmap at the bottom depicts the relative enrichment of the indicated functional groups in both classes. (f) Bar graphs depicting the prediction performance of Metabokiller on the indicated unseen datasets.

**Supplementary Figure 3:**
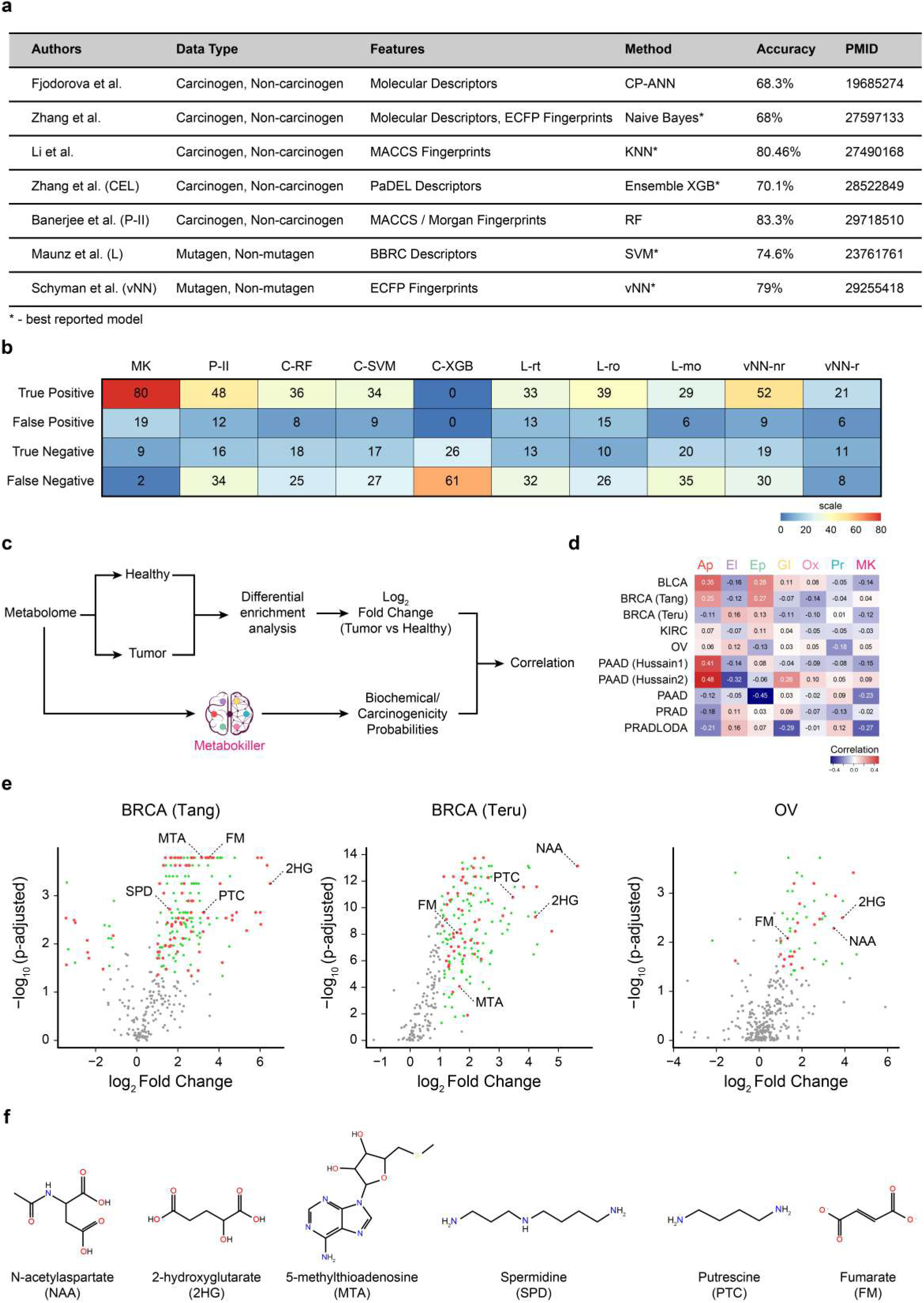
Metabokiller outperforms state-of-the-art methods for carcinogenicity prediction. **(a)** Table summarizing the source information and reported performance of the publicly available model for carcinogenicity or mutagenicity prediction. **(b)** Heatmap depicting the number of true positive (TP), false positive (FP), true negative (TN), and false negative (FN) predictions on the Independent Dataset (I.D.) for indicated methods/tools. **(c)** Schematic representation of the steps involved in processing pan-cancer metabolomics dataset. Of note, Pearson correlation was computed between log_2_ fold change (tumor vs healthy) and biochemical/carcinogenicity probabilities. **(d)** Heatmap detailing the correlation values further segregated based on cancer type. **(e)** Volcano plots depicting the differentially enriched/de-enriched metabolites in the indicated cancer datasets. Grey dots highlight the metabolites that do not qualify for the enrichment cutoff (log_2_ fold change ≥1 or ≤ -1, and p-value (adjusted) < 0.05), and green and red dots represent the metabolites that qualify for the enrichment cutoff and are predicted as non-carcinogenic and carcinogenic by Metabokiller respectively. **(f)** Structural information of some of the well-characterized oncometabolites reported in the literature and predicted by Metabokiller.

**Supplementary Figure 4:**
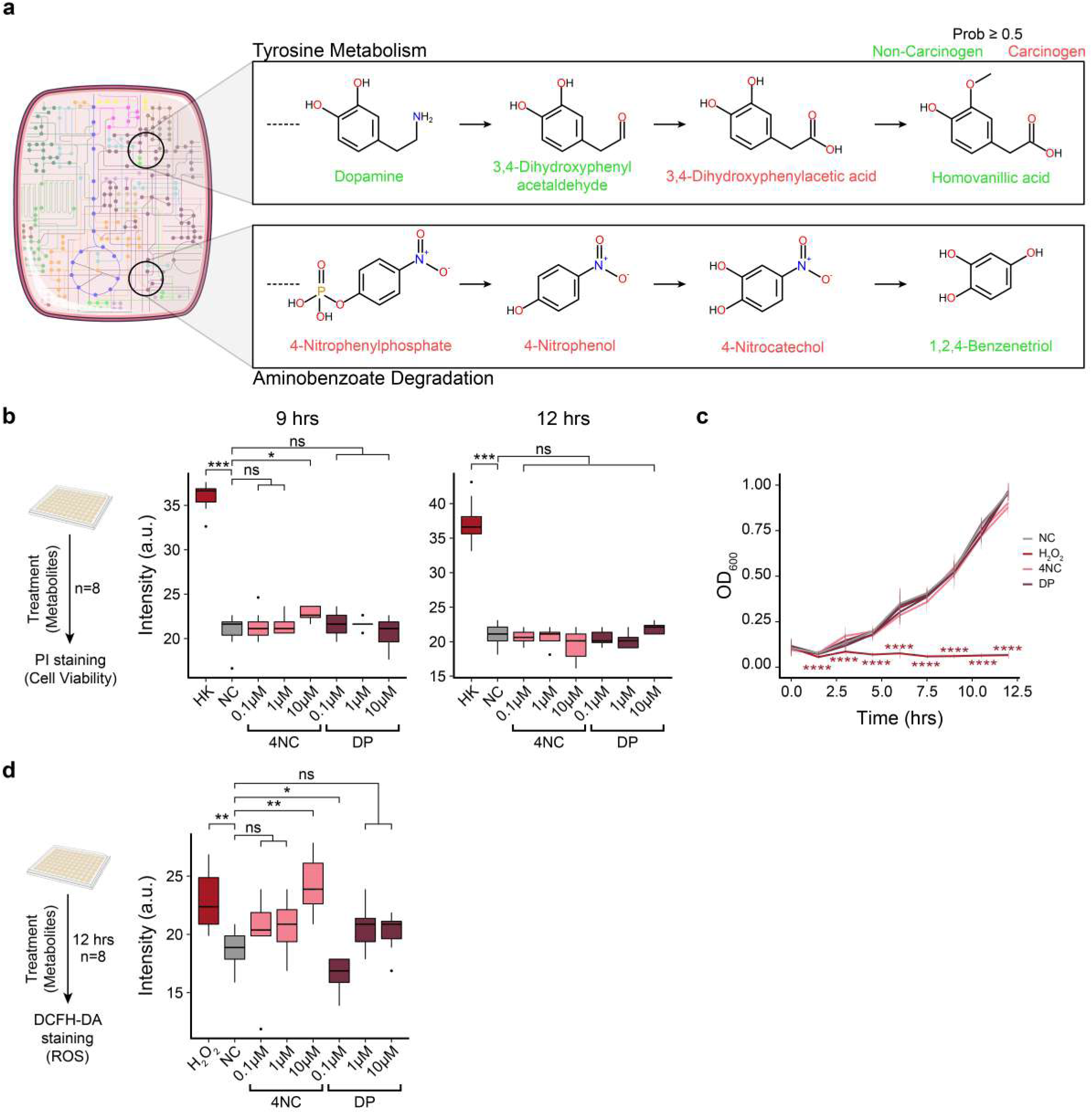
High synergy between Metabokiller predictions and experimental validations. **(a)** Schematic representation highlighting the predicted-carcinogenic metabolic intermediates of the tyrosine metabolism pathway and aminobenzoate degradation pathway. **(b)** Box plots depicting the fluorescence intensity of propidium iodide staining indicating cell viability in the indicated conditions. Of note, heat-killed (HK) yeast cells were used as a positive control. Mann–Whitney U test was used to compute statistical significance between the test conditions and the negative control. **(c)** Growth curve profiles of the treated and untreated wild-type yeast during transient exposure with the indicated conditions. The Student’s t-test was used to compute statistical significance between the positive (H_2_O_2_ treated yeast cells) and negative control (untreated yeast cells). **(d)** Box plot depicting the results of reactive oxygen species (ROS) levels inferred using DCFH-DA dye-based assay in the indicated conditions. Of note, ROS levels were measured 12 hours post-incubation. Mann–Whitney U test was used to compute statistical significance between the test conditions and the negative control. Notably, hydrogen peroxide (H_2_O_2_) treated yeast cells were used as a positive control.

**Supplementary Figure 5:**
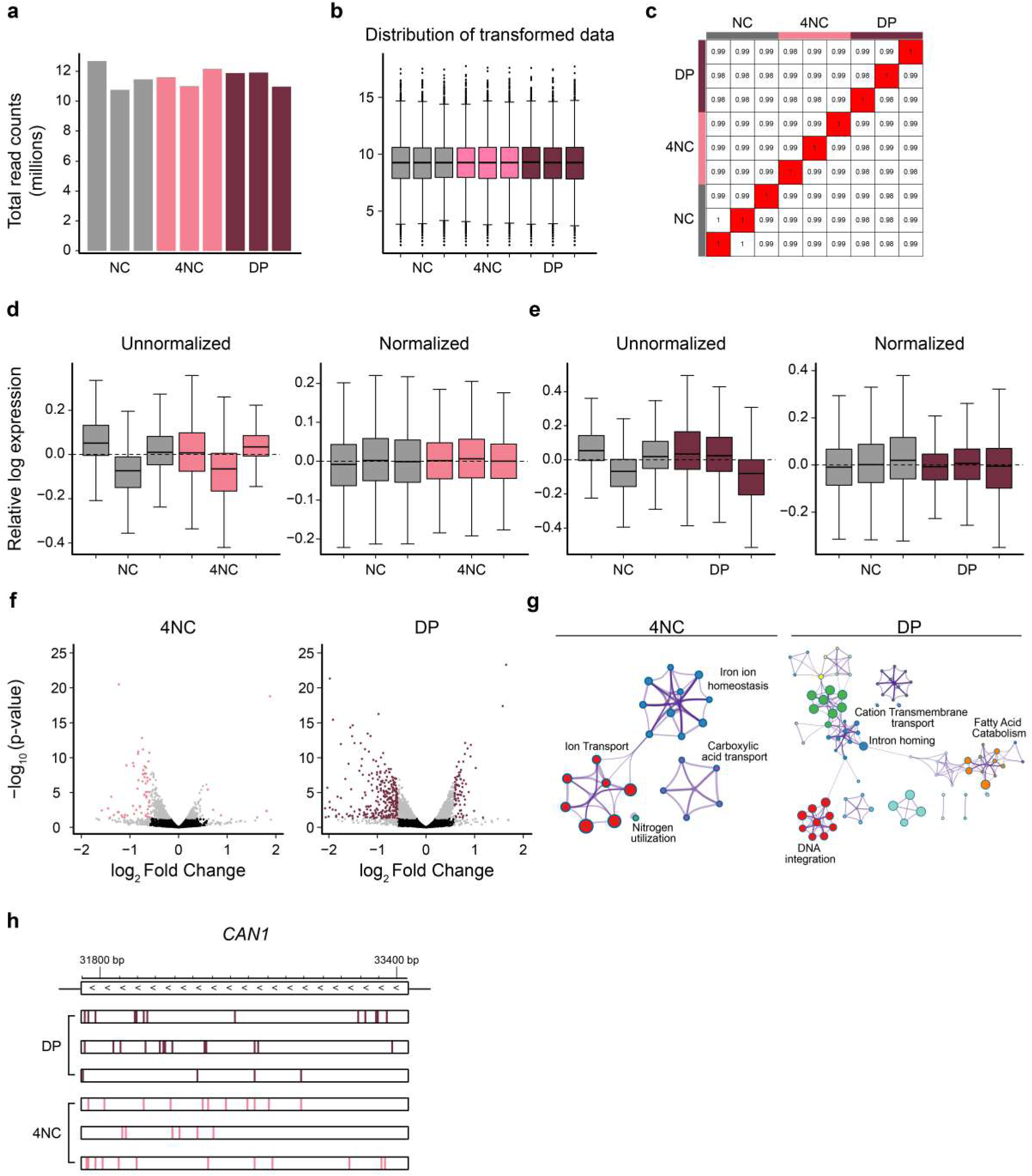
Deep RNA sequencing revealed the prominent molecular and biological processes affected in 4-Nitrocatechol and 3,4-Dihydroxyphenylacetic acid treated yeast cells. **(a)** Bar plots depicting the total read counts (in millions) of the indicated RNA sequencing samples. **(b)** Box plot representing the distribution of the transformed read count data in the indicated conditions. **(c)** Correlation plot showing the relationship between the individual RNA sequencing samples. Of note, 75% of the normalized and transformed data was used for the correlation analysis. **(d-e)** Box plots depicting the relative log expression of the individual replicates of the indicated conditions before and after upper quantile normalization. **(f)** Volcano plot indicating the differentially expressed genes between the treated (metabolite treatment) and untreated conditions. **(g)** Metascape-based Functional Gene Ontology analysis identified the involvement of differentially expressed genes in the indicated prominent biological processes. **(h)** Schematic representation depicting the genomic alterations in the *CAN1* gene in the indicated replicates.

## Notes

https://github.com/the-ahuja-lab/Metabokiller

https://doi.org/10.5281/zenodo.6477021

https://pypi.org/project/Metabokiller/

